# Vaccine Protection Against the SARS-CoV-2 Omicron Variant in Macaques

**DOI:** 10.1101/2022.02.06.479285

**Authors:** Abishek Chandrashekar, Jingyou Yu, Katherine McMahan, Catherine Jacob-Dolan, Jinyan Liu, Xuan He, David Hope, Tochi Anioke, Julia Barrett, Benjamin Chung, Nicole P. Hachmann, Michelle Lifton, Jessica Miller, Olivia Powers, Michaela Sciacca, Daniel Sellers, Mazuba Siamatu, Nehalee Surve, Haley VanWyk, Huahua Wan, Cindy Wu, Laurent Pessaint, Daniel Valentin, Alex Van Ry, Jeanne Muench, Mona Boursiquot, Anthony Cook, Jason Velasco, Elyse Teow, Adrianus C.M. Boon, Mehul S. Suthar, Neharika Jain, Amanda J. Martinot, Mark G. Lewis, Hanne Andersen, Dan H. Barouch

**Author notes:** Co-first author (Abishek Chandrashekar, Jingyou Yu, Katherine McMahan, Catherine Jacob-Dolan, Jinyan Liu, Xuan He, David Hope). Corresponding author: Dan H. Barouch, M.D., Ph.D., Center for Virology and Vaccine Research, 330 Brookline Avenue, E/CLS-1043, Boston, MA 02115.

## Abstract

**Background:** The rapid spread of the SARS-CoV-2 Omicron (B.1.1.529) variant, including in highly vaccinated populations, has raised important questions about the efficacy of current vaccines. Immune correlates of vaccine protection against Omicron are not known.

**Methods:** 30 cynomolgus macaques were immunized with homologous and heterologous prime-boost regimens with the mRNA-based BNT162b2 vaccine and the adenovirus vector-based Ad26.COV2.S vaccine. Following vaccination, animals were challenged with the SARS-CoV-2 Omicron variant by the intranasal and intratracheal routes.

**Results:** Omicron neutralizing antibodies were observed following the boost immunization and were higher in animals that received BNT162b2, whereas Omicron CD8+ T cell responses were higher in animals that received Ad26.COV2.S. Following Omicron challenge, sham controls showed more prolonged virus in nasal swabs than in bronchoalveolar lavage. Vaccinated macaques demonstrated rapid control of virus in bronchoalveolar lavage, and most vaccinated animals also controlled virus in nasal swabs, showing that current vaccines provide substantial protection against Omicron in this model. However, vaccinated animals that had moderate levels of Omicron neutralizing antibodies but negligible Omicron CD8+ T cell responses failed to control virus in the upper respiratory tract. Virologic control correlated with both antibody and T cell responses.

**Conclusions:** BNT162b2 and Ad26.COV2.S provided robust protection against high-dose challenge with the SARS-CoV-2 Omicron variant in macaques. Protection against this highly mutated SARS-CoV-2 variant correlated with both humoral and cellular immune responses.

## INTRODUCTION

The highly mutated SARS-CoV-2 Omicron variant has been shown to evade neutralizing antibody (NAb) responses induced by current vaccines, although a third immunization augments Omicron-specific NAb responses^1–5^. In contrast, T cell responses induced by current vaccines have been reported to be highly cross-reactive to SARS-CoV-2 variants including Omicron^6–8^.

Recent clinical effectiveness studies have shown that the mRNA-based BNT162b2 vaccine^9^ and the adenovirus vector-based Ad26.COV2.S vaccine^10^ provided 70% and 85% protection, respectively, against hospitalization with Omicron in South Africa^11, 12^. In this study, we evaluated the immunogenicity and protective efficacy of BNT162b2 and Ad26.COV2.S, including homologous and heterologous boost regimens, against SARS-CoV-2 Omicron challenge in nonhuman primates.

## METHODS

### Animals, Vaccines, and Challenge Stock

30 adult male and female cynomolgus macaques ages 4-12 years old were randomly allocated to 5 experimental groups (N=6/group; **Fig. S1**). Groups of animals were primed with either two immunizations of 30 μg BNT162b2 at weeks 0 and 3 or a single immunization of 5×10^10^ vp Ad26.COV2.S at week 0. At week 14, animals were boosted with either 30 μg BNT162b2 or 5×10^10^ vp Ad26.COV2.S. Clinical vaccines were obtained from pharmacies by the NIH SAVE Consortium. At week 19, all animals were challenged with 10^6^ PFU SARS-CoV-2 Omicron by the intranasal and intratracheal routes in a total volume of 2 mls. This challenge stock was generated in VeroE6-TMPRSS2 cells and had a titer of 2.3×10^9^ TCID50/ml and 2.5×10^7^ PFU/ml in VeroE6-TMPRSS2 cells and was fully sequenced (EPI_ISL_7171744; Mehul Suthar, Emory University). Following challenge, viral loads were assessed in bronchoalveolar lavage (BAL) and nasal swab (NS) samples by RT-PCR for E subgenomic RNA (sgRNA), and infectious virus titers were quantitated by TCID50 assays. Animals were sacrificed on day 9 or 10 following challenge. Immunologic and virologic assays were performed blinded. All animals were housed at Bioqual, Inc. (Rockville, MD). All animal studies were conducted in compliance with all relevant local, state, and federal regulations and were approved by the Bioqual Institutional Animal Care and Use Committee (IACUC).

### Pseudovirus neutralizing antibody assay

The SARS-CoV-2 pseudoviruses expressing a luciferase reporter gene were used to measure pseudovirus neutralizing antibodies^13^. In brief, the packaging construct psPAX2 (AIDS Resource and Reagent Program), luciferase reporter plasmid pLenti-CMV Puro-Luc (Addgene) and spike protein expressing pcDNA3.1-SARS-CoV-2 SΔCT were co-transfected into HEK293T cells (ATCC CRL_3216) with lipofectamine 2000 (ThermoFisher Scientific). Pseudoviruses of SARS-CoV-2 variants were generated by using WA1/2020 strain (Wuhan/WIV04/2019, GISAID accession ID: EPI_ISL_402124), B.1.617.2 (Delta, GISAID accession ID: EPI_ISL_2020950), or B.1.1.529 (Omicron, GISAID ID: EPI_ISL_7358094.2). The supernatants containing the pseudotype viruses were collected 48h after transfection; pseudotype viruses were purified by filtration with 0.45-μm filter. To determine the neutralization activity of human serum, HEK293T-hACE2 cells were seeded in 96-well tissue culture plates at a density of 2.0 × 10^4^ cells per well overnight. Three-fold serial dilutions of heat-inactivated serum samples were prepared and mixed with 50 μl of pseudovirus. The mixture was incubated at 37 °C for 1 h before adding to HEK293T-hACE2 cells. After 48 h, cells were lysed in Steady-Glo Luciferase Assay (Promega) according to the manufacturer’s instructions. SARS-CoV-2 neutralization titers were defined as the sample dilution at which a 50% reduction (NT50) in relative light units was observed relative to the average of the virus control wells.

### Enzyme-linked immunosorbent assay (ELISA)

SARS-CoV-2 spike receptor-binding domain (RBD)-specific binding antibodies in serum were assessed by ELISA. 96-well plates were coated with 1 μg/mL of similarly produced SARS-CoV-2 WA1/2020, B.1.617.2 (Delta), B.1.351 (Beta), or B.1.1.529 (Omicron) RBD protein in 1× Dulbecco phosphate-buffered saline (DPBS) and incubated at 4 °C overnight. Assay performance was similar for these four RBD proteins. After incubation, plates were washed once with wash buffer (0.05% Tween 20 in 1× DPBS) and blocked with 350 μL of casein block solution per well for 2 to 3 hours at room temperature. Following incubation, block solution was discarded and plates were blotted dry. Serial dilutions of heat-inactivated serum diluted in Casein block were added to wells, and plates were incubated for 1 hour at room temperature, prior to 3 more washes and a 1-hour incubation with a 1μg/mL dilution of anti–macaque IgG horseradish peroxidase (HRP) (Nonhuman Primate Reagent Resource) at room temperature in the dark. Plates were washed 3 times, and 100 μL of SeraCare KPL TMB SureBlue Start solution was added to each well; plate development was halted by adding 100 μL of SeraCare KPL TMB Stop solution per well. The absorbance at 450 nm was recorded with a VersaMax microplate reader (Molecular Devices). For each sample, the ELISA end point titer was calculated using a 4-parameter logistic curve fit to calculate the reciprocal serum dilution that yields an absorbance value of 0.2. Interpolated end point titers were reported.

### Electrochemiluminescence assay (ECLA)

ECLA plates (MesoScale Discovery SARS-CoV-2 IgG, Panels 22, 23) were designed and produced with up to 10 antigen spots in each well, including Spike and RBD from multiple SARS-CoV-2 variants^14^. The plates were blocked with 50 uL of Blocker A (1% BSA in distilled water) solution for at least 30 minutes at room temperature shaking at 700 rpm with a digital microplate shaker. During blocking the serum was diluted to 1:5,000 or 1:50,000 in Diluent 100. The calibrator curve was prepared by diluting the calibrator mixture from MSD 1:10 in Diluent 100 and then preparing a 7-step 4-fold dilution series plus a blank containing only Diluent 100. The plates were then washed 3 times with 150 μL of Wash Buffer (0.5% Tween in 1x PBS), blotted dry, and 50 μL of the diluted samples and calibration curve were added in duplicate to the plates and set to shake at 700 rpm at room temperature for at least 2 h. The plates were again washed 3 times and 50 μL of SULFO-Tagged anti-Human IgG detection antibody diluted to 1x in Diluent 100 was added to each well and incubated shaking at 700 rpm at room temperature for at least 1 h. Plates were then washed 3 times and 150 μL of MSD GOLD Read Buffer B was added to each well and the plates were read immediately after on a MESO QuickPlex SQ 120 machine. MSD titers for each sample was reported as Relative Light Units (RLU) which were calculated as Sample RLU minus Blank RLU and then fit using a logarithmic fit to the standard curve. The upper limit of detection was defined as 2×10^6 RLU for each assay and the signal for samples which exceeded this value at 1:5,000 serum dilution was run again at 1:50,000 and the fitted RLU was multiplied by 10 before reporting. The lower limit of detection was defined as 1 RLU and an RLU value of 100 was defined to be positive for each assay.

### Intracellular cytokine staining (ICS) assay

CD4+ and CD8+ T cell responses were quantitated by pooled peptide-stimulated intracellular cytokine staining (ICS) assays. Peptide pools were 16 amino acid peptides overlapping by 11 amino acids spanning the SARS-CoV-2 WA1/2020, B.1.617.2 (Delta), or B.1.1.529 (Omicron; GISAID ID: EPI_ISL_7358094.2) Spike proteins (21^st^ Century Biochemicals). 10^6^ peripheral blood mononuclear cells well were re-suspended in 100 µL of R10 media supplemented with CD49d monoclonal antibody (1 µg/mL) and CD28 monoclonal antibody (1 µg/mL). Each sample was assessed with mock (100 µL of R10 plus 0.5% DMSO; background control), peptides (2 µg/mL), and/or 10 pg/mL phorbol myristate acetate (PMA) and 1 µg/mL ionomycin (Sigma-Aldrich) (100µL; positive control) and incubated at 37°C for 1 h. After incubation, 0.25 µL of GolgiStop and 0.25 µL of GolgiPlug in 50 µL of R10 was added to each well and incubated at 37°C for 8 h and then held at 4°C overnight. The next day, the cells were washed twice with DPBS, stained with aqua live/dead dye for 10 mins and then stained with predetermined titers of monoclonal antibodies against CD279 (clone EH12.1, BB700), CD38 (clone OKT10, PE), CD28 (clone 28.2, PE CY5), CD4 (clone L200, BV510), CD95 (clone DX2, BUV737), CD8 (clone SK1, BUV805) for 30 min. Cells were then washed twice with 2% FBS/DPBS buffer and incubated for 15 min with 200 µL of BD CytoFix/CytoPerm Fixation/Permeabilization solution. Cells were washed twice with 1X Perm Wash buffer (BD Perm/WashTM Buffer 10X in the CytoFix/CytoPerm Fixation/ Permeabilization kit diluted with MilliQ water and passed through 0.22µm filter) and stained with intracellularly with monoclonal antibodies against Ki67 (clone B56, FITC), CD69 (clone TP1.55.3, ECD), IL10 (clone JES3-9D7, PE CY7), IL13 (clone JES10-5A2, BV421), TNF-α (clone Mab11, BV650), IL4 (clone MP4-25D2, BV711), IFN-γ (clone B27; BUV395), CD45 (clone D058-1283, BUV615), IL2 (clone MQ1-17H12, APC), CD3 (clone SP34.2, Alexa 700)for 30 min. Cells were washed twice with 1X Perm Wash buffer and fixed with 250µL of freshly prepared 1.5% formaldehyde. Fixed cells were transferred to 96-well round bottom plate and analyzed by BD FACSymphony™ system. Data were analyzed using FlowJo v9.9.

### B cell immunophenotyping

PBMCs or inguinal LN cells were stained with Aqua live/dead dye for 20 minutes, washed with 2% FBS/DPBS buffer, and cells were suspended in 2% FBS/DPBS buffer with Fc Block (BD Biosciences) for 10 minutes^15^. After blocking, samples were stained with monoclonal antibodies against CD45 (clone D058-1283, brilliant ultraviolet (BUV) 805), CD3 (clone SP34.2, allophycocyanin (APC)-Cy7), CD7 (clone M-T701, Alexa Fluor700), CD123 (clone 6H6, Alexa Fluor 700), CD11c (clone 3.9, Alexa Fluor 700), CD19 (clone J3-119, phycoerythrin (PE)), CD20 (clone 2H7, PE-Cy5), IgD (IA6-2, PE), IgG (clone G18-145, BUV737), IgM (clone G20-127, BUV395), CD80 (clone L307.4, brilliant violet (BV) 786), CD95 (clone DX2, BV711), CD27 (clone M-T271, BUV563), CD21 (clone B-ly4, BV605), CD14 (clone M5E2, BV570). Samples were also stained with SARS-CoV-2 antigens, including biotinylated SARS-CoV-2 (WA1/2020) RBD proteins (Sino Biological), SARS-CoV-2 (WA1/2020) RBD proteins (Sino Biological) labeled with fluorescein isothiocyanate (FITC), SARS-CoV-2 (B.1.1.529) RBD proteins (Sino Biological) labeled with APC and DyLight 405. Staining was done at 4 °C for 30 minutes. After staining, cells were washed twice with 2% FBS/DPBS buffer, followed by incubation with BV650 streptavidin (BD Pharmingen) for 10 minutes, then washed twice with 2% FBS/DPBS buffer. For intracellular staining, cells were permeabilized using Caltag Fix & Perm (Thermo Fisher Scientific), then stained with monoclonal antibodies against Ki67 (clone B56, peridinin chlorophyll protein (PerCP)-Cy5.5) and Bcl6 (clone K112-91, PE-CF594). After staining, cells were washed and fixed by 2% paraformaldehyde. All data were acquired on a BD FACSymphony flow cytometer. Subsequent analyses were performed using FlowJo software (BD Bioscience, v.9.9.6). For analyses, in singlet gate, dead cells were excluded by Aqua dye and CD45 was used as a positive inclusion gate for all leukocytes. Within class-switched memory B cell populations, gated as CD20+IgG+CD27+IgM-CD3-CD14-CD11c-CD123-CD7-, SARS-CoV-2 WA1/2020 RBD-specific B cells were identified as double positive for SARS-CoV-2 (WA1/2020) RBD labeled with different fluorescent probes, and SARS-CoV-2 (B.1.1.529) RBD-specific B cells were identified as double positive for SARS-CoV-2 (B.1.1.529) RBD proteins labeled with different fluorescent probes. Within GC B cells gated as CD20+ IgD- IgG+ Ki67+ Bcl6+, SARS-CoV-2 RBD-specific GC B cells were identified as double positive for SARS-CoV-2 RBD with different fluorescent probes.

### Subgenomic RT-PCR assay

SARS-CoV-2 E gene subgenomic RNA (sgRNA) was assessed by RT-PCR using primers and probes as previously described^13^. A standard was generated by first synthesizing a gene fragment of the subgenomic E gene. The gene fragment was subsequently cloned into a pcDNA3.1+ expression plasmid using restriction site cloning (Integrated DNA Technologies). The insert was in vitro transcribed to RNA using the AmpliCap-Max T7 High Yield Message Maker Kit (CellScript). Log dilutions of the standard were prepared for RT-PCR assays ranging from 1×1010 copies to 1×10-1 copies. Viral loads were quantified from bronchoalveolar lavage (BAL) fluid and nasal swabs (NS). RNA extraction was performed on a QIAcube HT using the IndiSpin QIAcube HT Pathogen Kit according to manufacturer’s specifications (Qiagen). The standard dilutions and extracted RNA samples were reverse transcribed using SuperScript VILO Master Mix (Invitrogen) following the cycling conditions described by the manufacturer. A Taqman custom gene expression assay (Thermo Fisher Scientific) was designed using the sequences targeting the E gene sgRNA. The sequences for the custom assay were as follows, forward primer, sgLeadCoV2.Fwd: CGATCTCTTGTAGATCTGTTCTC, E_Sarbeco_R: ATATTGCAGCAGTACGCACACA, E_Sarbeco_P1 (probe): VIC-ACACTAGCCATCCTTACTGCGCTTCG-MGBNFQ. Reactions were carried out in duplicate for samples and standards on the QuantStudio 6 and 7 Flex Real-Time PCR Systems (Applied Biosystems) with the thermal cycling conditions: initial denaturation at 95°C for 20 seconds, then 45 cycles of 95°C for 1 second and 60°C for 20 seconds. Standard curves were used to calculate subgenomic RNA copies per ml or per swab. The quantitative assay sensitivity was determined as 50 copies per ml or per swab.

### TCID50 assay

Vero-TMPRSS2 cells (obtained from A. Creanga) were plated at 25,000 cells per well in DMEM with 10% FBS and gentamicin, and the cultures were incubated at 37 °C, 5.0% CO2. Medium was aspirated and replaced with 180 μl of DMEM with 2% FBS and gentamicin. Serial dilution of samples as well as positive (virus stock of known infectious titre) and negative (medium only) controls were included in each assay. The plates are incubated at 37 °C, 5.0% CO2 for 4 days. Cell monolayers were visually inspected for cytopathic effect. The TCID50 was calculated using the Read–Muench formula.

### Histopathology and Immunohistochemistry

Lungs from SARS CoV-2 WA1/2020 and Omicron infected macaques were evaluated on day 2 following challenge by histopathology^16^. At the time of fixation, lungs were suffused with 10% formalin to expand the alveoli. All tissues were fixed in 10% formalin and blocks sectioned at 5 μm. Slides were incubated for 30–60 min at 65°C then deparaffinized in xylene and rehydrated through a series of graded ethanol to distilled water. Sections were stained with hematoxylin and eosin. For SARS-N immunohistochemistry, heat-induced epitope retrieval was performed using a pressure cooker on steam setting for 25 min in citrate buffer (Thermo Fisher Scientific, AP-9003–500), followed by treatment with 3% hydrogen peroxide. Slides were then rinsed in distilled water and protein blocked (Biocare, BE965H) for 15 min followed by rinses in 1× PBS. Primary mouse anti-SARS-CoV-nucleoprotein antibody (Sinobiological; 40143-MM05) at 1:1000, was applied for 60 min, followed by mouse Mach-2 HRP-Polymer (Biocare) for 30 min and then counterstained with hematoxylin followed by bluing using 0.25% ammonia water. Staining was performed using a Biocare intelliPATH autostainer. Blinded evaluation and histopathologic scoring of eight representative lung lobes from cranial, middle and caudal, left and right lungs from each monkey was performed by a board-certified veterinary pathologist (AJM).

### Statistical analysis

Descriptive statistics and logistic regression were performed using GraphPad Prism 8.4.3, (GraphPad Software, San Diego, California). Immunologic data were generated in duplicate and were compared by two-sided Mann-Whitney tests. Correlations were assessed by two-sided Spearman rank-correlation tests. P values less than 0.05 were considered significant.

## RESULTS

### Humoral Immune Responses

We immunized 30 adult cynomolgus macaques with homologous and heterologous regimens with BNT162b2 and Ad26.COV2.S or sham vaccine (N=6/group; **Fig. S1**). Groups of animals were primed with either two immunizations of 30 μg BNT162b2 at weeks 0 and 3 or a single immunization of 5×10^10^ vp Ad26.COV2.S at week 0. At week 14, animals received a homologous or heterologous boost with 30 μg BNT162b2 or 5×10^10^ vp Ad26.COV2.S.

NAb responses were evaluated by luciferase-based pseudovirus neutralizing antibody assays^13^. Vaccine-matched WA1/2020 NAbs were induced in all animals after the priming immunization at week 8 and were 13.3-fold higher in the BNT162b2 primed animals compared with the Ad26.COV2.S primed animals. The WA1/2020 NAb titers in the BNT162b2 vaccinated groups declined more than 10-fold by week 14 (**Fig. 1A**), consistent with immune kinetics following BNT162b2 vaccination in humans^17, 18^. Omicron-specific NAbs were low in all groups prior to the boost. At week 18 after the homologous and heterologous boosts, median WA1/2020 NAb titers were 19,901, 15,451, 7,461, 2,215, and <20 in the BNTx3, BNTx2/Ad26, Ad26/BNT, Ad26×2, and sham groups, respectively. Median Omicron NAb titers at week 18 were 1,901, 650, 810, 168, and <20, respectively, reflecting a 9-23 fold reduction compared with WA1/2020 NAb titers (**Fig. 1A**).

**Figure 1.**
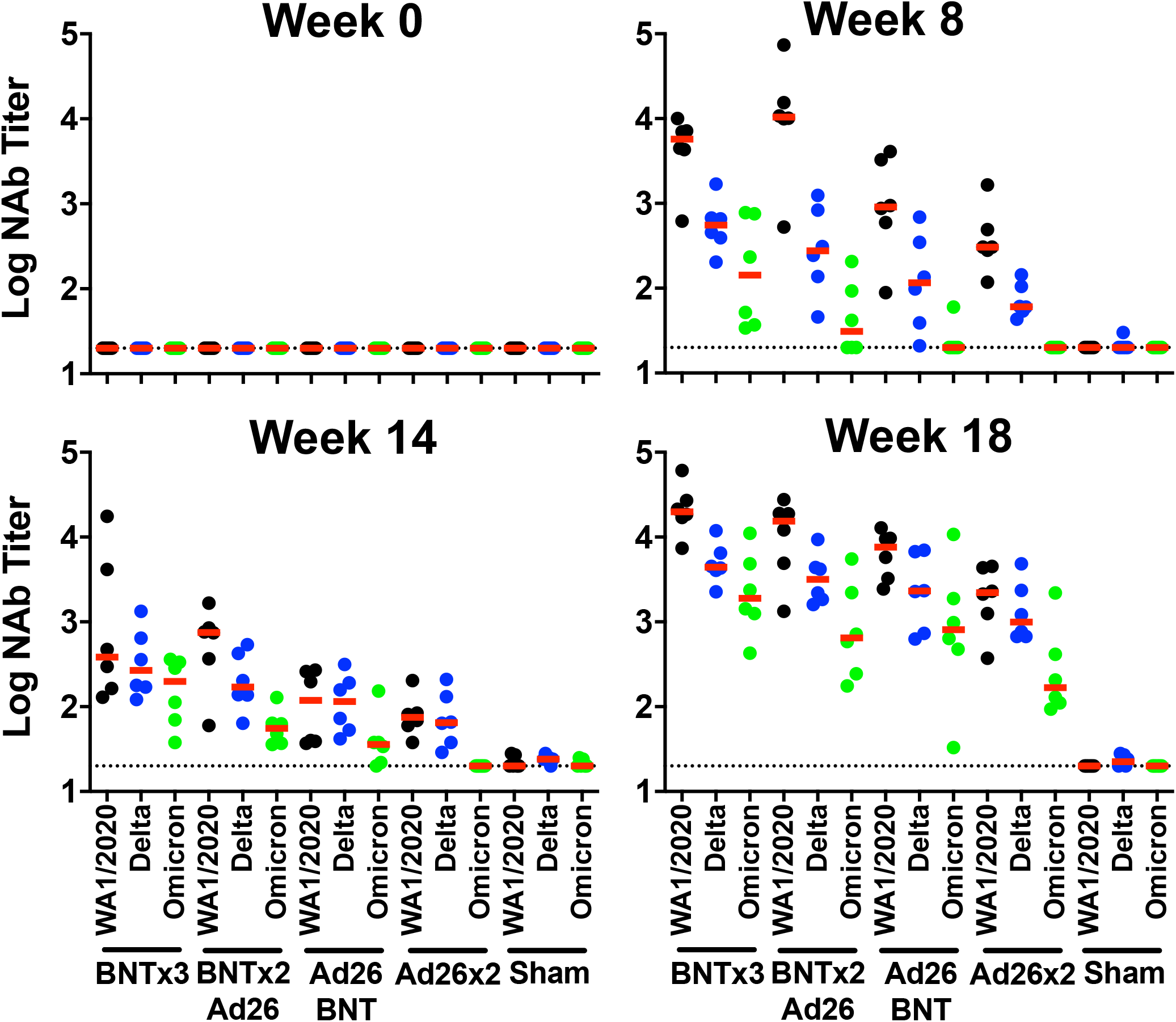
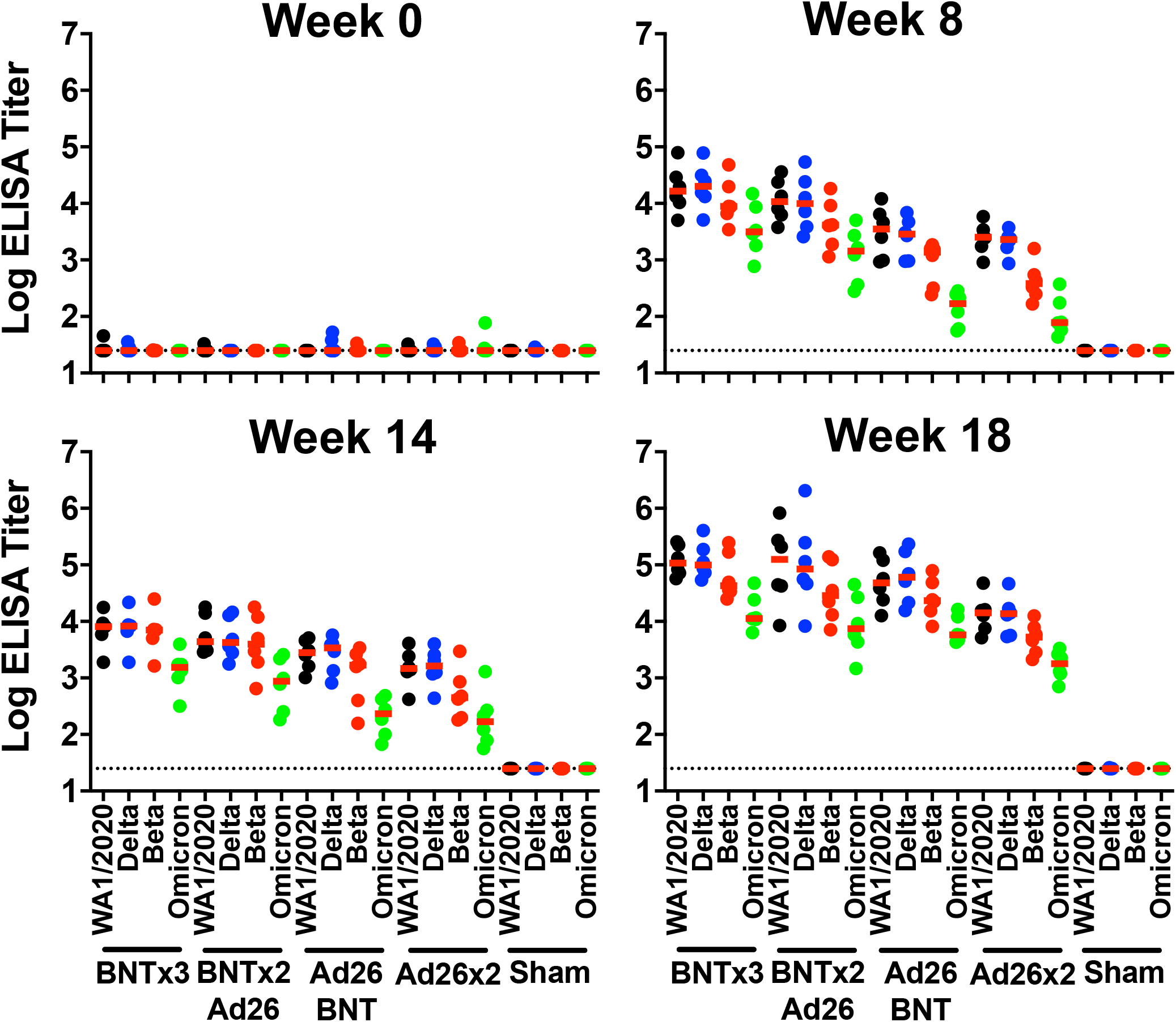
Humoral immune responses following vaccination. Antibody responses at weeks 0 (baseline), 8 (post-prime), 14 (pre-boost), and 18 (post-boost) following vaccination with BNTx3, BNTx2/Ad26, Ad26/BNT, Ad26×2, or sham (N=30; N=6/group). **A,** Neutralizing antibody (NAb) titers by a luciferase-based pseudovirus neutralization assay. **B**, Receptor binding domain (RBD)-specific binding antibody titers by ELISA. Responses were measured against the SARS-CoV-2 WA1/2020 (black), B.1.617.2 (Delta; blue), B.1.351 (Beta; red), and B.1.1.529 (Omicron; green) variants. Dotted lines represent limits of quantitation. Medians (red bars) are shown.

Receptor-binding domain (RBD) specific binding antibodies were assessed by ELISA. At week 18, median WA1/2020 ELISA titers were 107,705, 125,694, 60,634, 14,193, and <25 in the BNTx3, BNTx2/Ad26, Ad26/BNT, Ad26×2, and sham groups, respectively. Median Omicron ELISA titers were 11,333, 7,452, 5,805, 1,783, and <25, respectively, reflecting a 8-17 fold reduction compared with WA1/2020 ELISA titers (**Fig. 1B**). Similar trends were observed in Spike- and RBD-specific binding assays using the Meso-Scale Discovery electrochemiluminescence assay (ECLA)^14^ (**Figs. S2, S3**). These data show that homologous and heterologous boosts substantially increased antibody responses in all groups, although Omicron antibody responses remained approximately 10-fold lower than WA1.2020 antibody responses.

### Cellular Immune Responses

We next assessed Spike-specific CD8+ and CD4+ T cell responses by multiparameter flow cytometry. At week 14 prior to the boost, WA1/2020 Spike-specific IFN-γ CD8+ T cell responses were 13.1-fold higher in the Ad26.COV2.S primed animals compared with the BNT162b2 primed animals (**Fig. 2A**), consistent with cellular immune data in humans^17, 19, 20^. In contrast, WA1/2020 Spike-specific IFN-γ CD4+ T cell responses were comparable across groups (**Fig. 2B**). Moreover, for both CD8+ and CD4+ T cell responses, Omicron responses were similar to WA1/2020 responses, indicative of substantial cross-reactivity of T cell responses^6–8, 21^. At week 16 after the homologous and heterologous boosts, median Omicron Spike-specific IFN-γ CD8+ T cell responses were 0.012%, 0.023%, 0.034%, 0.031%, and 0.004% in the BNTx3, BNTx2/Ad26, Ad26/BNT, Ad26×2, and sham groups, respectively (**Fig. 2A**). Median Omicron Spike-specific IFN-γ CD4+ T cell responses were 0.150%, 0.088%, 0.081%, 0.028%, and 0.001% in the BNTx3, BNTx2/Ad26, Ad26/BNT, Ad26×2, and sham groups, respectively (**Fig. 2B**).

**Figure 2.**
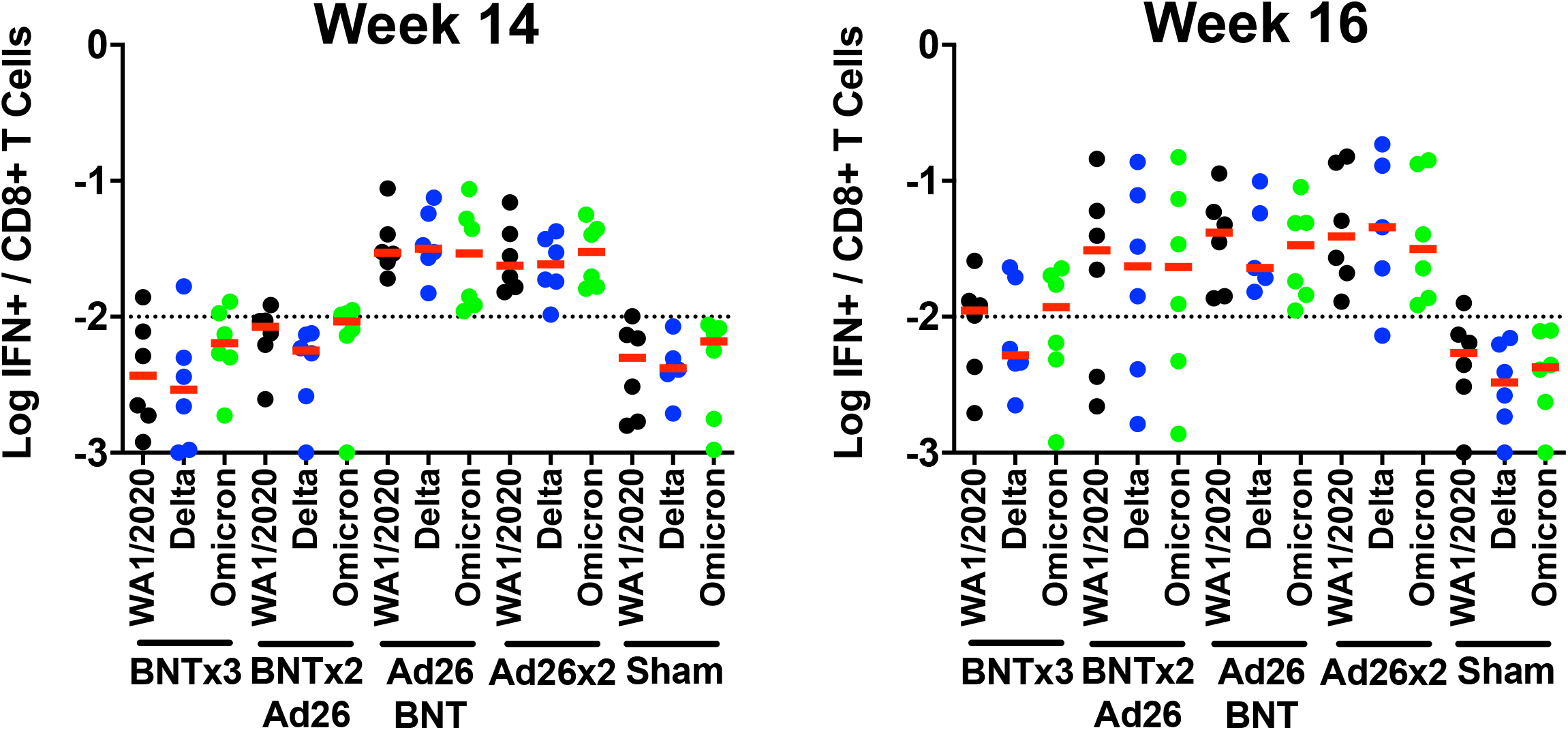
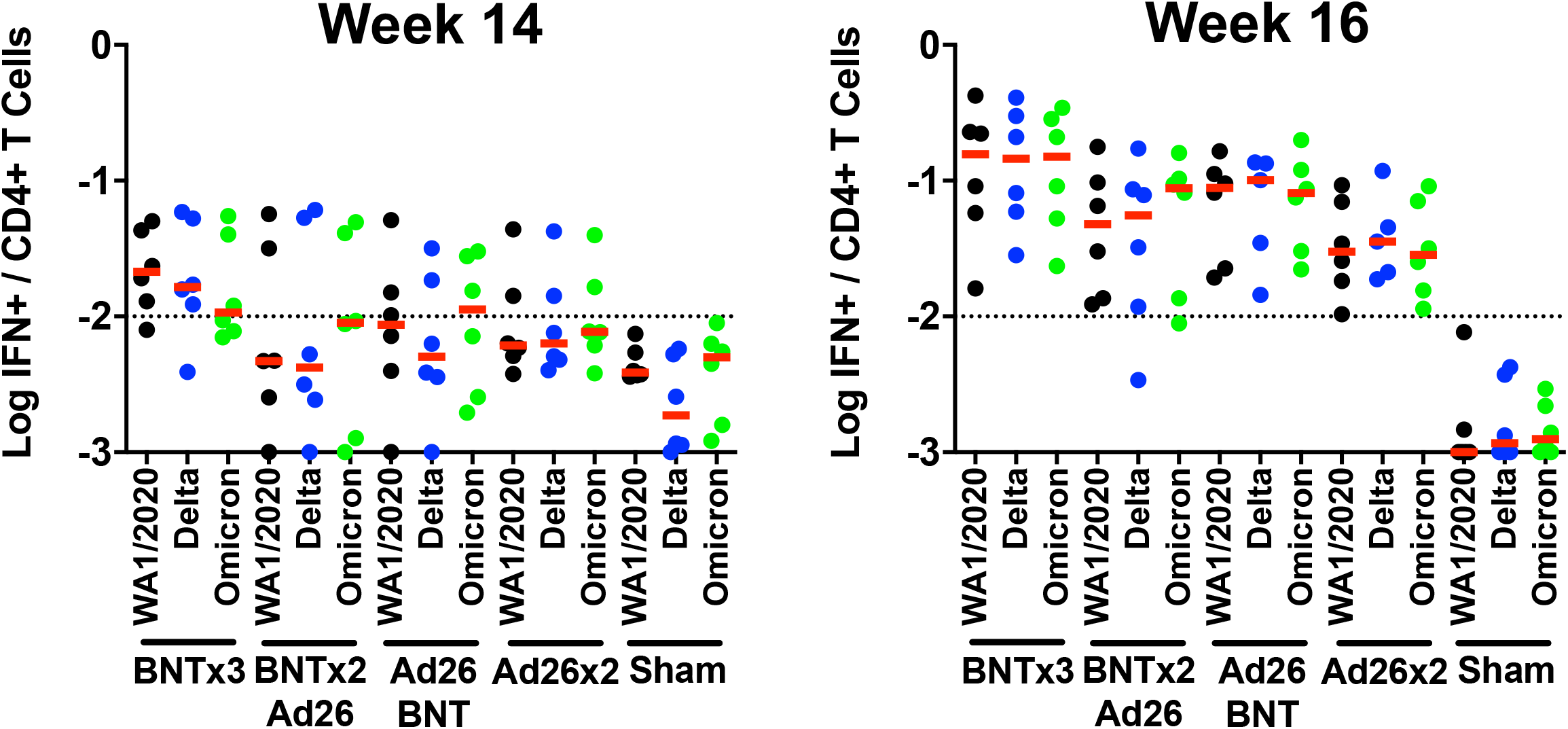
Cellular immune responses following vaccination. T cell responses at weeks 14 (pre-boost) and 18 (post-boost) following vaccination with BNTx3, BNTx2/Ad26, Ad26/BNT, Ad26×2, or sham (N=30; N=6/group). Pooled peptide Spike-specific IFN-γ (**A**) CD8+ T cell responses and (**B**) CD4+ T cell responses by intracellular cytokine staining assays. Responses were measured against the SARS-CoV-2 WA1/2020 (black), B.1.617.2 (Delta; blue), and B.1.1.529 (Omicron; green) variants. Dotted lines represent limits of quantitation. Medians (red bars) are shown.

We also assessed memory IgG+ B cells in peripheral blood as well as germinal center CD20+IgD-IgG+Ki67+Bcl6+ B cells in lymph nodes at week 16 by multiparameter flow cytometry. Omicron RBD-specific memory B cells and germinal center B cells were induced at comparable levels in all vaccinated groups (**Fig. S4**). Peripheral Omicron RBD-specific memory B cells correlated with lymph node Omicron RBD-specific germinal center B cells (R=0.6543, P=0.0002, two-sided Spearman rank-correlation test) and serum Omicron NAb titers (R=0.5602, P=0.0019, two-sided Spearman rank-correlation test) at week 16 (**Fig. S5**).

### Protective Efficacy Following SARS-CoV-2 Omicron Challenge

At week 19, all animals were challenged with 10^6^ PFU SARS-CoV-2 Omicron by the intranasal and intratracheal routes. This challenge stock was generated in VeroE6-TMPRSS2 cells and had a titer of 2.3×10^9^ TCID50/ml and 2.5×10^7^ PFU/ml in VeroE6-TMPRSS2 cells, and the sequence of the challenge stock was fully verified (EPI_ISL_7171744; Mehul Suthar, Emory University). Following challenge, viral loads were assessed in bronchoalveolar lavage (BAL) and nasal swab (NS) samples by RT-PCR for E subgenomic RNA (sgRNA)^22, 23^, and infectious virus titers were quantitated by TCID50 assays.

Sham controls showed high median viral loads of 5.70 (range 4.84-7.36) log sgRNA copies/ml in BAL on day 2, and these levels declined substantially by day 7 to median levels of 2.82 (range 1.78-4.10) log sgRNA copies/ml (**Fig. 3A**). Nearly all vaccinated animals demonstrated breakthrough infection in BAL, but viral loads were substantially lower in vaccinated animals compared with sham controls on day 2 and mostly resolved by day 4 (**Fig. 3A**). In NS, sham controls showed median virus levels of 4.06 (range 3.05-4.59) log sgRNA copies/ml on day 2, and these levels only declined minimally by day 7 to median levels of 3.85 (range 3.50-4.49) log sgRNA copies/ml (**Fig. 3B**). All vaccinated animals showed breakthrough infection in NS, but viral loads resolved in most vaccinated animals by day 4, with the exception of 2 animals in the BNTx3 group and 2 animals in the BNTx2/Ad26 group that showed persistent high levels of virus in NS through day 7, which was comparable with sham controls (**Fig. 3B**).

**Figure 3.**
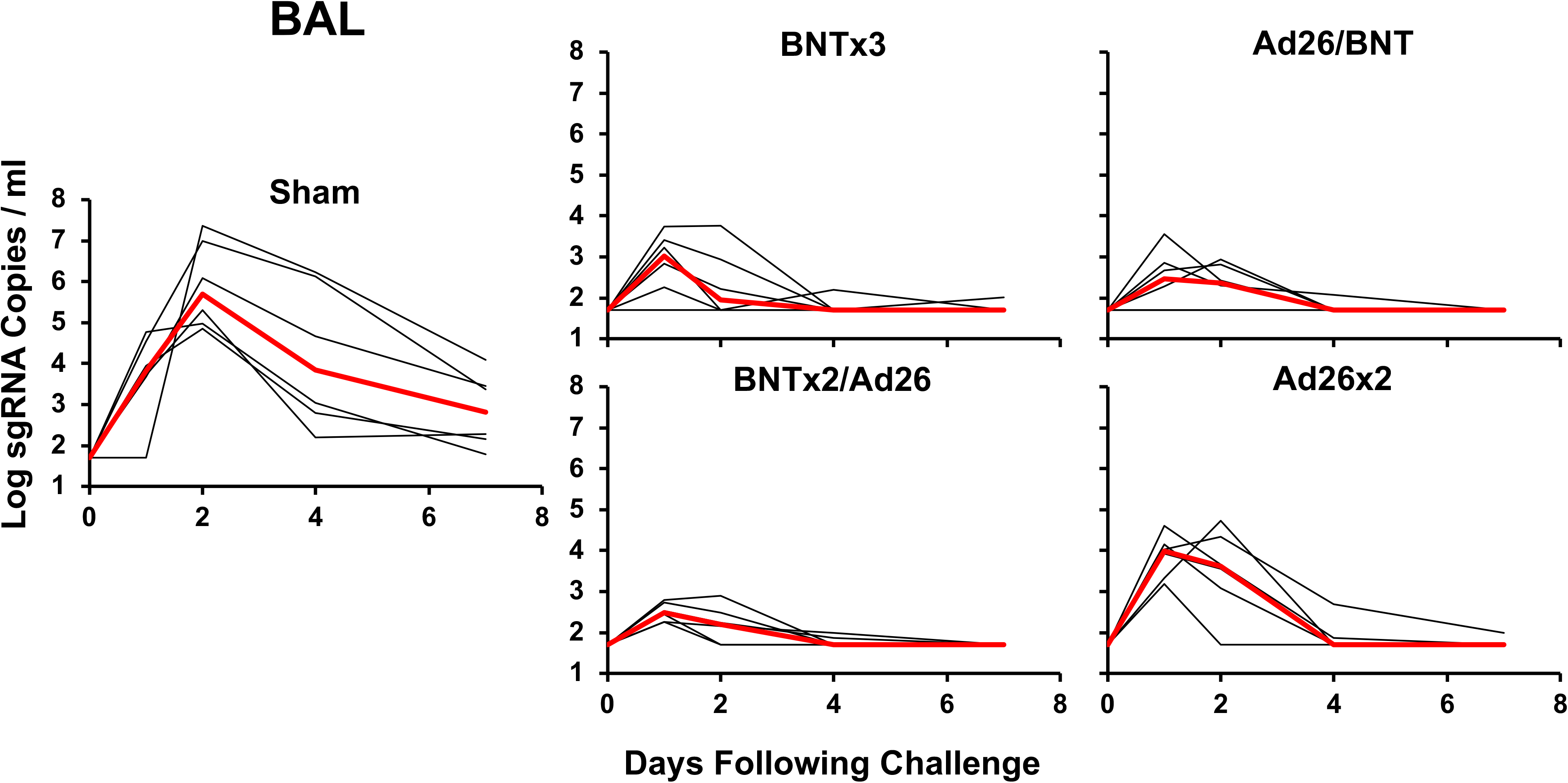
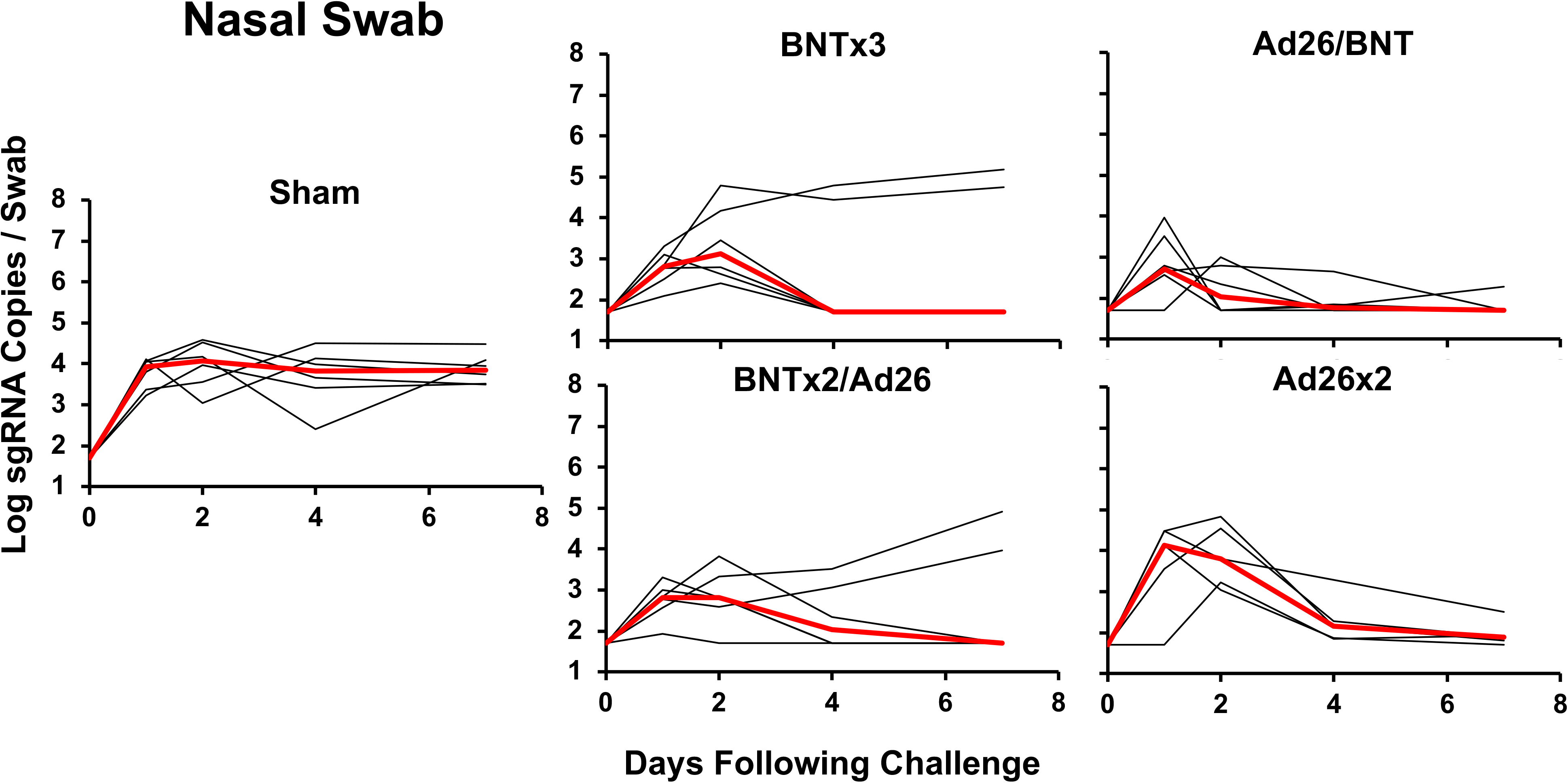
Viral loads following SARS-CoV-2 Omicron challenge. **A,** Log subgenomic RNA (sgRNA) copies/ml in bronchoalveolar lavage (BAL) following SARS-CoV-2 Omicron challenge. **B,** Log subgenomic RNA (sgRNA) copies/swab in nasal swabs (NS) following SARS-CoV-2 Omicron challenge. Medians (red lines) are shown.

Median log peak viral loads in BAL were reduced by 2.68-, 3.21-, 2.87-, and 1.46-fold in the BNTx3, BNTx2/Ad26, Ad26/BNT, and Ad26×2 groups, respectively, compared with sham controls (P=0.0022, P=0.0022, P=0.0022, P=0.0022, respectively, two-tailed Mann-Whitney tests, **Fig. 4A**). Median log day 4 viral loads in BAL were also significantly reduced to undetectable levels in all groups compared with sham controls (P=0.0022, P=0.0022, P=0.0022, P=0.0043, respectively, two-tailed Mann-Whitney tests, **Fig. 4A**). Median log peak viral loads in NS were only reduced in the heterologous BNTx2/Ad26 and Ad26/BNT groups compared with sham controls (P=0.0043 and P=0.0043, respectively, two-tailed Mann-Whitney tests, **Fig. 4B**). Median log day 4 viral loads in NS were reduced in the BNTx2/Ad26, Ad26/BNT, and Ad26×2 groups compared with sham controls (P=0.0152, P=0.0043, and P=0.0087, respectively, two-tailed Mann-Whitney tests, **Fig. 4B**). Consistent with the RT-PCR viral load data, vaccinated animals also showed substantial log reductions of infectious virus titers compared with sham controls in BAL and NS by TCID50 assays on day 2 (**Fig. S6**).

**Figure 4.**
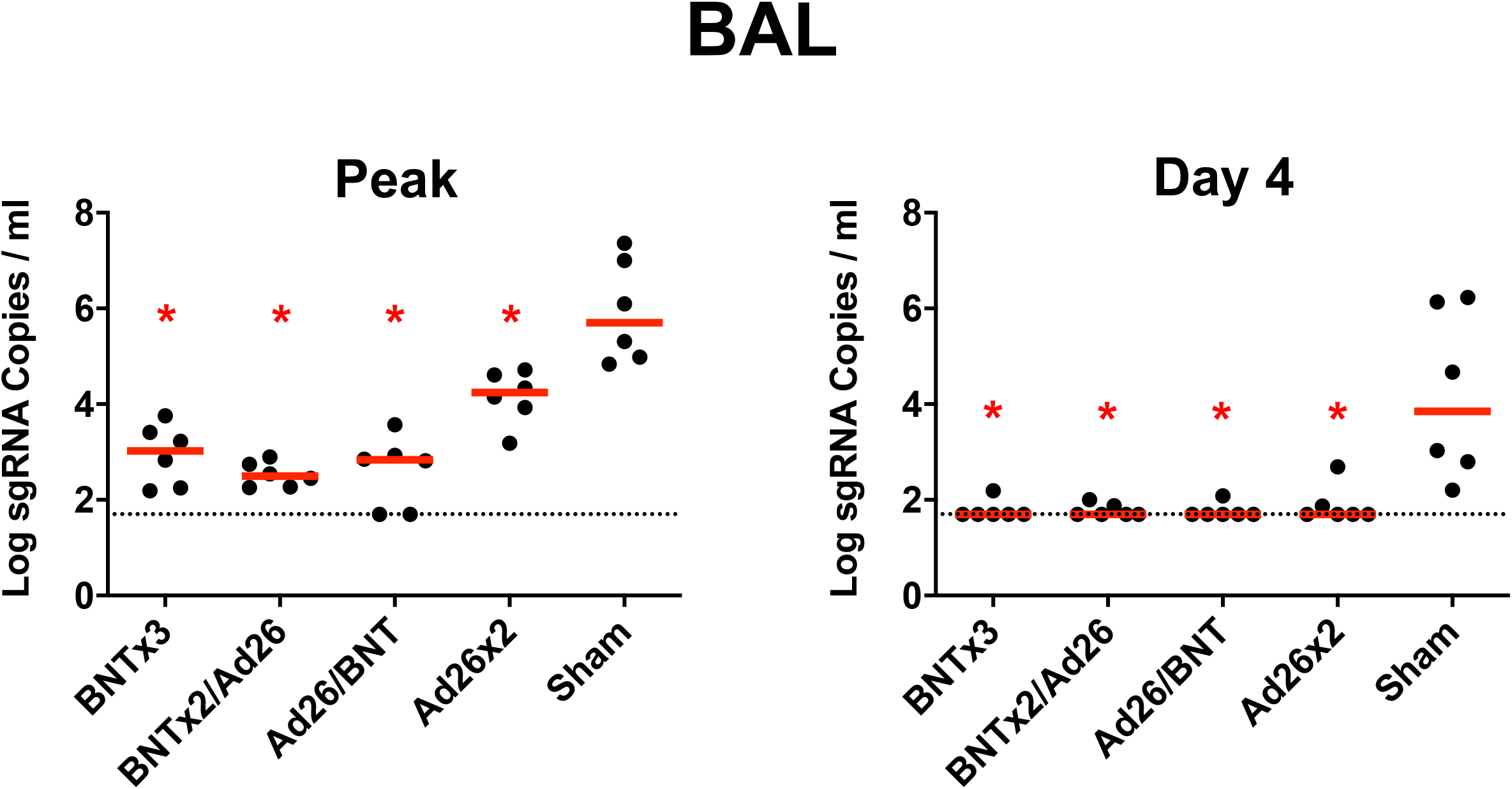
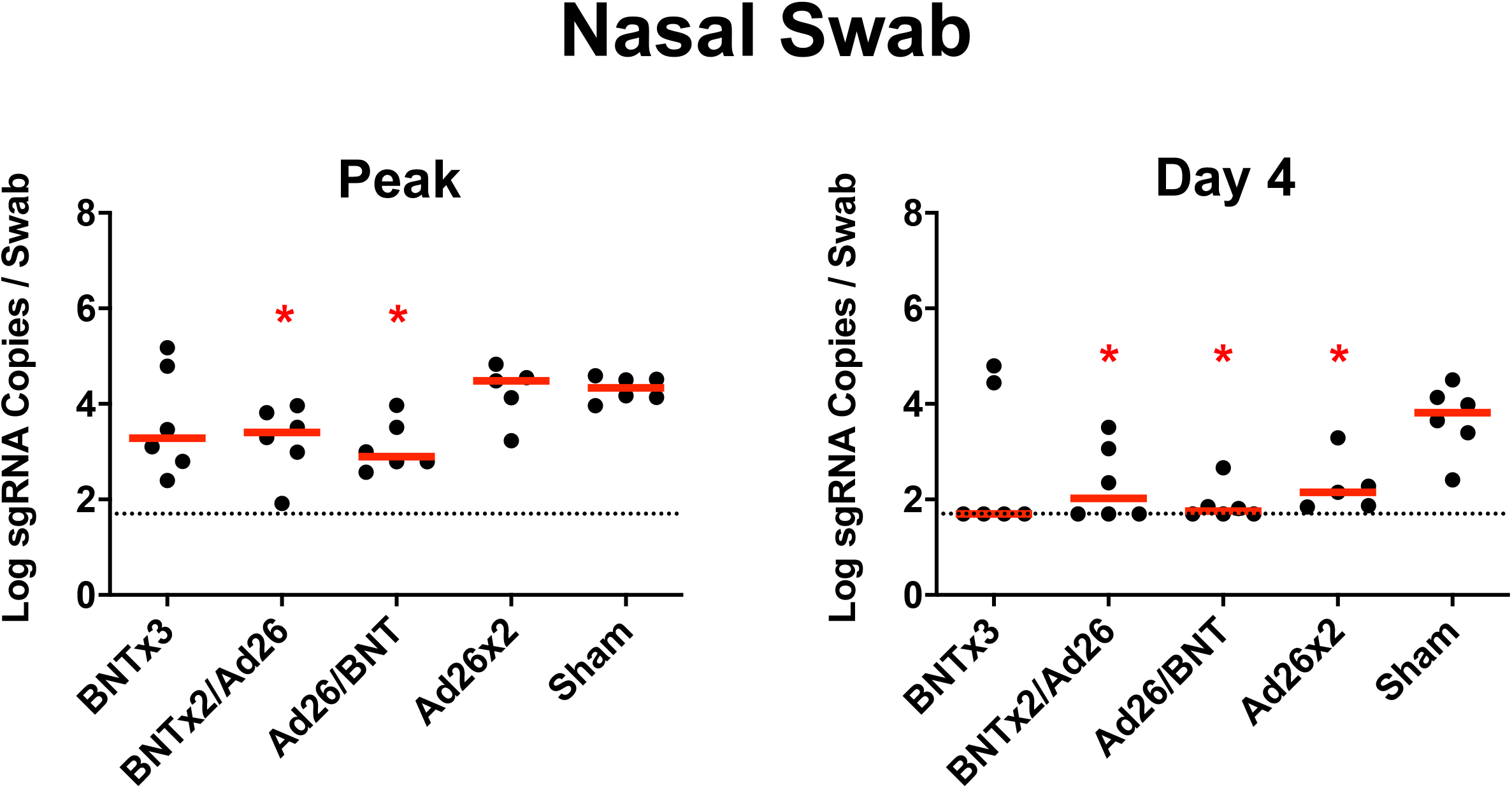
Comparison of peak and day 4 viral loads. **A,** Log subgenomic RNA (sgRNA) copies/ml in bronchoalveolar lavage (BAL) at peak and on day 4 following SARS-CoV-2 Omicron challenge. **B,** Log subgenomic RNA (sgRNA) copies/swab in nasal swabs (NS) at peak and on day 4 following SARS-CoV-2 Omicron challenge. Dotted lines represent limits of quantitation. Medians (red bars) are shown. Vaccinated groups were compared with the sham controls by two-sided Mann-Whitney tests. *, P<0.05.

### Correlates of Protection

We evaluated the immunologic profiles of the 4 vaccinated animals that failed to control viral replication in NS following challenge. These animals had moderate Omicron-specific NAb titers (586-1,434) but negligible Omicron-specific CD8+ T cell responses (0.001-0.006%) prior to challenge (red dots, **Fig. S7**). These 4 vaccinated animals and the 6 sham controls fell into a region of immunologic space defined by low to moderate Omicron NAbs and low Omicron CD8+ T cell responses (**Fig. 5**), suggesting that failure of virologic control following Omicron challenge was associated with simultaneously low humoral and cellular immunity to the challenge virus. In contrast, animals with a low NAb titer but a high CD8+ T cell response, or a high NAb titer but a low CD8+ T cell response, demonstrated rapid virologic control following challenge (red arrows, **Fig. 5**).

**Figure 5.**
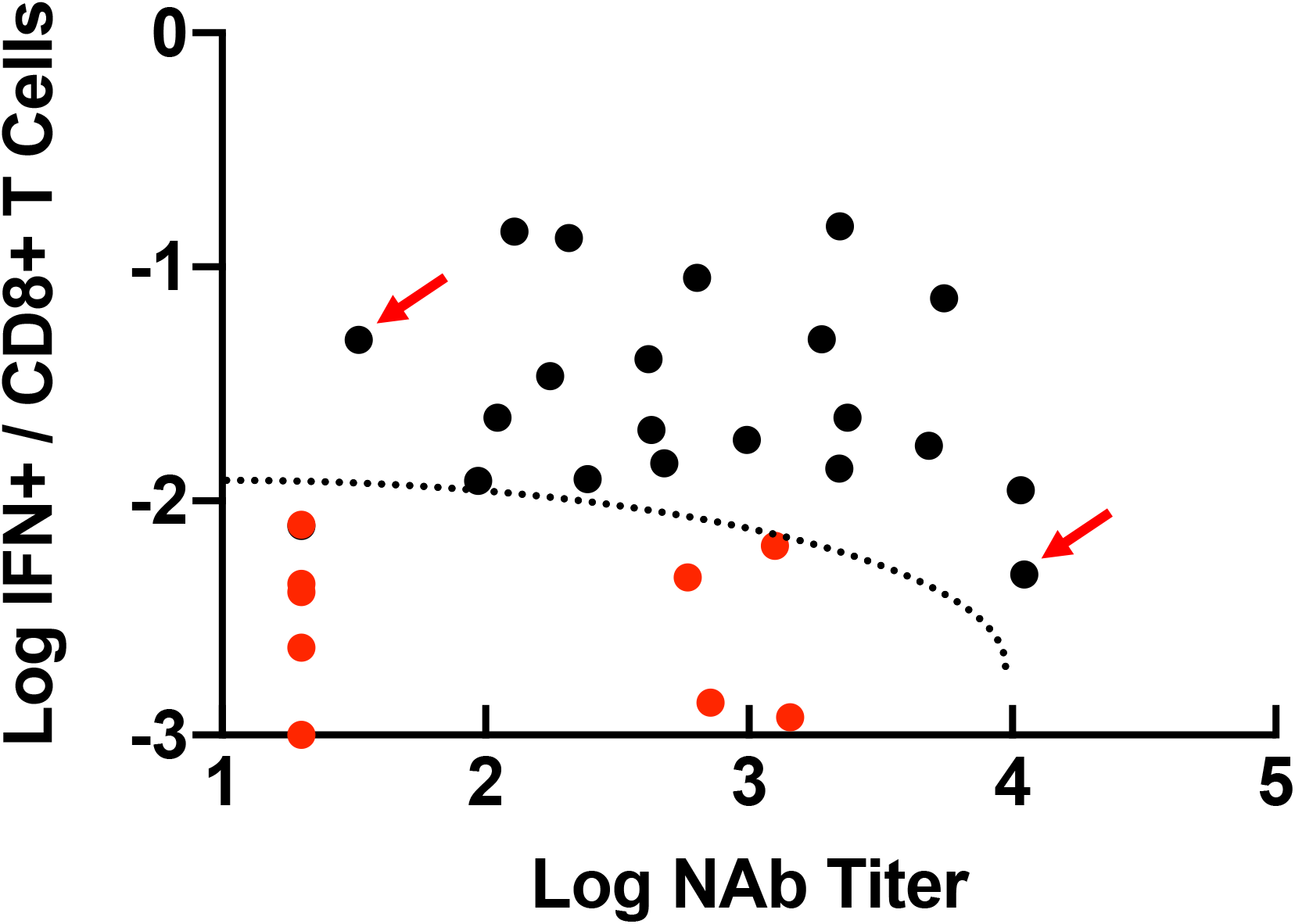
Immunologic space defined by Omicron NAb titer and Omicron CD8+ T cell responses. Plot of all 30 animals by their post-boost Omicron NAb titer and Omicron CD8+ T cell responses. Red dots represent the 10 animals that failed to control virus by day 7 in NS (6 controls, 4 vaccinated animals). The dotted line represents the region of immunologic space, defined post-hoc, which was associated with failure of virologic control. Red arrows show representative animals with low NAb titers but high CD8+ T cell responses, or high NAb titers but low CD8+ cell responses, which showed rapid virologic control.

The different magnitudes of humoral and cellular immune responses prior to challenge and viral loads following challenge allowed for a detailed immune correlates analysis. NAb titers, ELISA titers, CD8+ T cell responses, and CD4+ T cell responses all inversely correlated with sgRNA copies/ml in BAL (**Fig. S8**) and NS (**Fig. S9**). Since NAb titers and CD8+ T cell responses were not substantially correlated with each other (**Fig. 5**), these data suggest that both humoral and cellular immunity independently contributed to virologic control following Omicron challenge.

### Histopathology and immunohistochemistry

Histopathology and immunohistochemistry on day 2 following infection of additional macaques with the SARS-CoV-2 Omicron variant demonstrated lymphoid hyperplasia in the submucosa and rare SARS CoV-2 positive ciliated epithelial cells in the nasopharynx (**Fig. S10A-C**). Interstitial inflammation, expansion of septae, syncytial formation, and endothelialitis were observed in the lung in Omicron infected animals (**Fig. S10D-K**). However, lung histopathology scores were lower in macaques infected with Omicron compared with WA1/2020^16^ (P=0.0054, two-tailed Mann-Whitney test; **Fig. S10L**).

## DISCUSSION

In this study, we demonstrate that the BNT162b2 and Ad26.COV2.S vaccines led to rapid virologic control in the upper and lower respiratory tracts following high dose, heterologous challenge with the SARS-CoV-2 Omicron variant in the majority of macaques. However, 4 vaccinated animals with moderate Omicron-specific NAb titers but negligible Omicron-specific CD8+ T cell responses failed to control viral replication in NS by day 7. These data suggest the importance of vaccine-elicited CD8+ T cell responses and indicate that both humoral and cellular immune responses likely contribute to protection against the highly mutated SARS-CoV-2 Omicron variant in macaques.

Correlates of vaccine protection against SARS-CoV-2 infection have to date largely focused on neutralizing antibody titers^24, 25^, although correlates of protection against severe disease may differ from correlates of protection against infection, and the potential importance of vaccine-elicited T cell responses may be greater for SARS-CoV-2 variants such as Omicron that largely escape NAb responses. In the present study, Omicron-specific NAbs were markedly lower than WA1/2020 NAbs, whereas Omicron-specific T cell responses were similar to WA1/2020 T cell responses, indicating substantial cross-reactivity of cellular immune responses against SARS-CoV-2 variants. Moreover, while BNT162b2 induced higher NAb responses than Ad26.COV2.S, Ad26.COV2.S induced higher CD8+ T cell responses than BNT162b2, consistent with human data^17, 19, 20^. The different immune profiles induced by these vaccines suggest possible advantages of heterologous prime-boost (“mix-and-match”) vaccine regimens for diversifying immune responses.

We observed that virus persisted longer in NS compared with BAL in sham controls following Omicron challenge, which differs from prior SARS-CoV-2 variants in macaques^15, 16,26–29^. Although the implications of this observation remain to be determined, prolonged duration of virus shedding in the upper respiratory tract, together with substantial escape from NAbs, may contribute to the high degree of transmissibility of the SARS-CoV-2 Omicron variant.

Recent studies have shown that BNT162b2 and Ad26.COV2.S provided 70% and 85% protection, respectively, against hospitalization with Omicron in South Africa^11, 12^, largely in the absence of Omicron-specific NAbs. These data suggest that immune parameters other than NAb responses likely contribute to protection against severe disease. We previously reported that CD8+ T cells contributed to protection against re-challenge with SARS-CoV-2 in convalescent macaques, particularly when antibody responses were suboptimal^30^. Taken together, our data suggest that protection against a highly mutated SARS-CoV-2 variant involves a combination of humoral and cellular immunity, and not antibody responses alone. Moreover, moderate levels of neutralizing antibodies without substantial CD8+ T cell responses may be insufficient for virologic control. These data have important implications for understanding vaccine immune correlates against highly mutated SARS-CoV-2 variants.

## Acknowledgements

The authors thank S. Ducat, T. Hayes, A. Goode, G. Romero, J. Yalley-Ogunro, M. Cabus, B. Narvaez, E. Bouffard, S. Gardner, M. Gebre, V. Giffin, and Y. Tian for generous advice, assistance, and reagents. We thank MesoScale Discovery for providing the ECLA kits.

## Author Contributions

This study was designed by DHB. Immunologic and virologic assays were led by AC, JL, KM, CJD, JL, XH, DH, TA, JB, BC, NH, ML, JM, MS, DS, MS, OP, HV, HW, and CW. Humoral immune responses were assessed by CJD, KM, and JY. Vaccines were provided by ACMB. Omicron challenge stock was provided by MSS. Animal work was led by LP, DV, AVR, JM, MB, AC, JV, ET, MGL, and HA. Pathology was led by NJ and AJM. The paper was written by DHB with the involvement of all co-authors.

## Data Sharing

All data are available in the manuscript or the supplementary material.

## Correspondence and Requests

Correspondence and requests for materials should be addressed to D.H.B.

## SUPPLEMENTARY FIGURE LEGENDS

**Figure S1.**
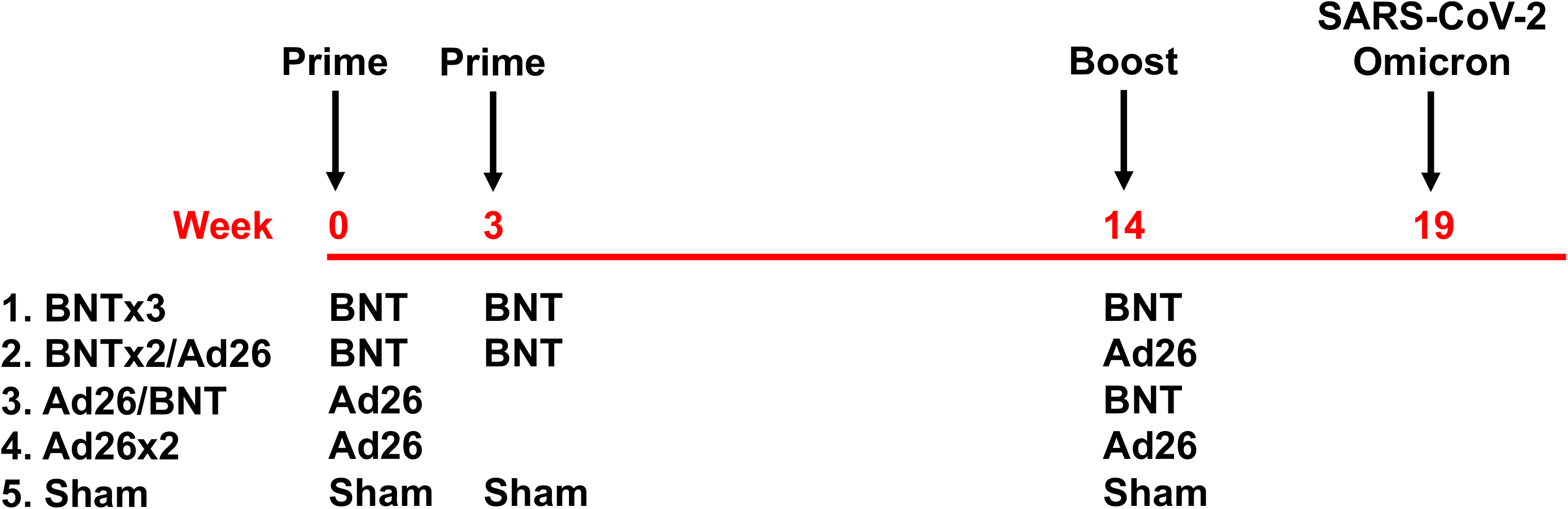
Study schema.

**Figure S2.**
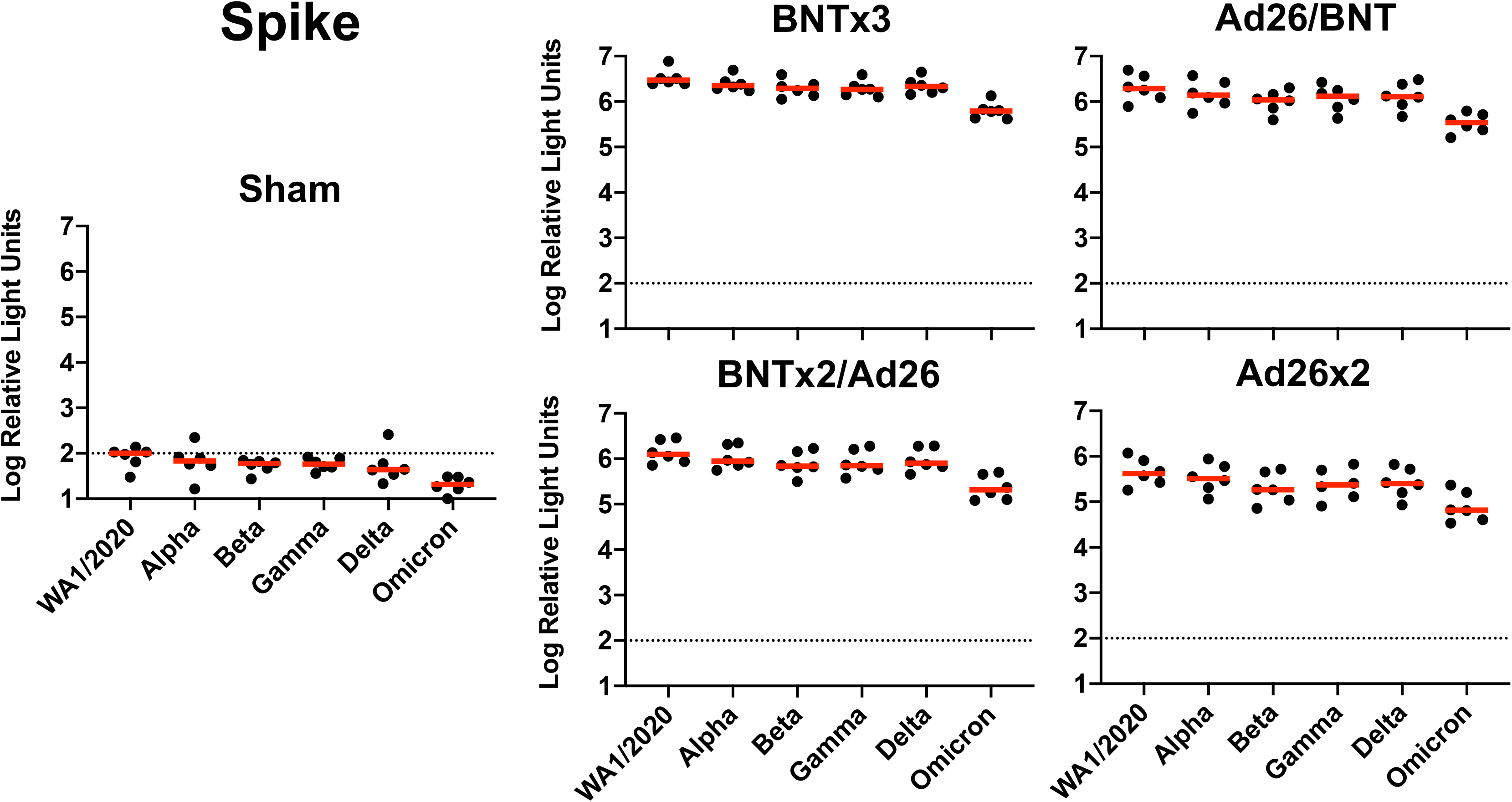
Spike-specific binding antibody responses following vaccination. Spike-specific antibody responses against multiple variants are shown at week 18 (post-boost) following vaccination with BNTx3, BNTx2/Ad26, Ad26/BNT, Ad26×2, or sham (N=30; N=6/group) with the Meso-Scale Discovery electrochemiluminescence assay (ECLA). Dotted lines represent limits of quantitation. Medians (red bars) are shown.

**Figure S3.**
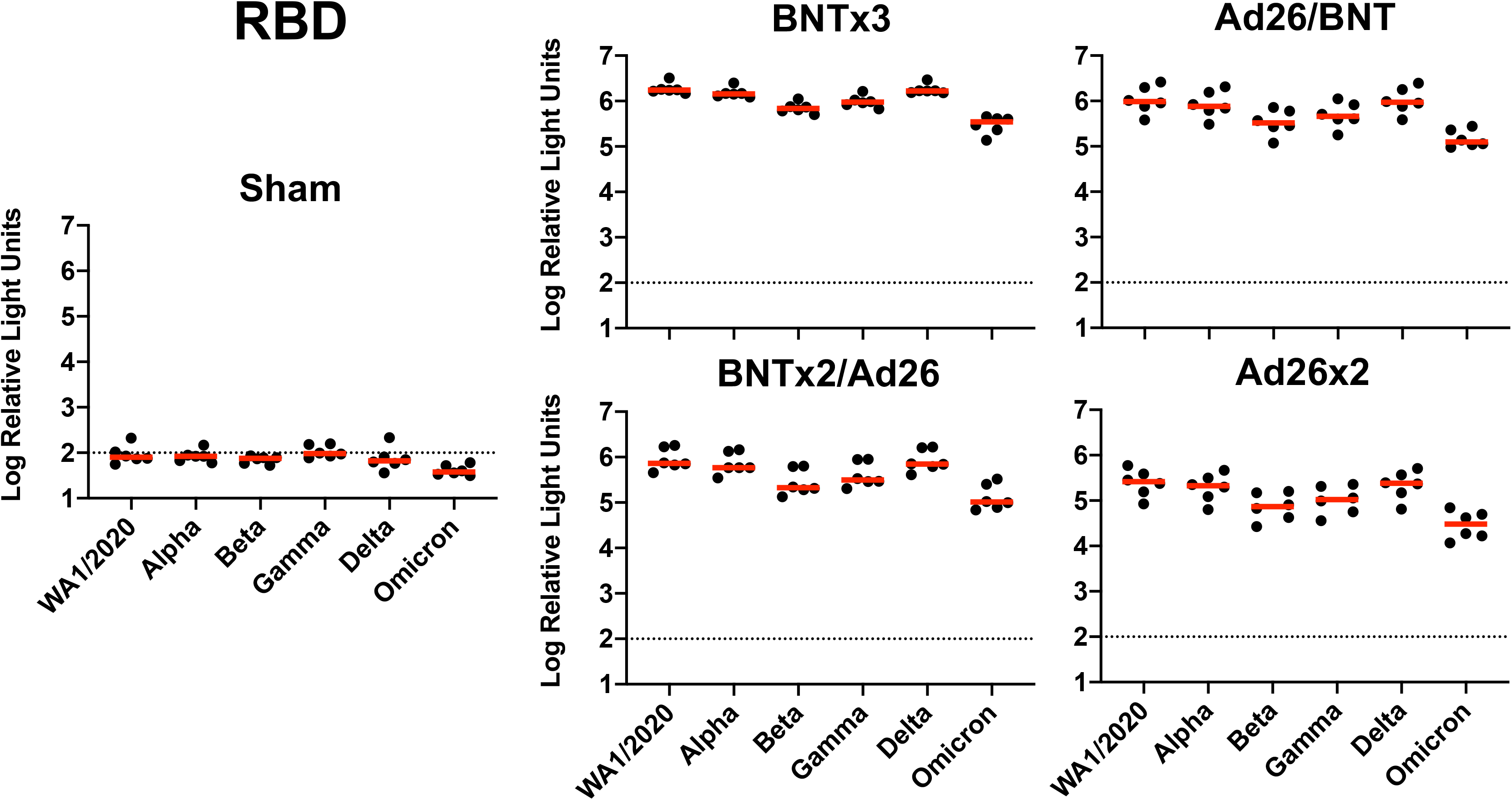
RBD-specific binding antibody responses following vaccination. RBD-specific antibody responses against multiple variants are shown at week 18 (post-boost) following vaccination with BNTx3, BNTx2/Ad26, Ad26/BNT, Ad26×2, or sham (N=30; N=6/group) with the Meso-Scale Discovery electrochemiluminescence assay (ECLA). Dotted lines represent limits of quantitation. Medians (red bars) are shown.

**Figure S4.**
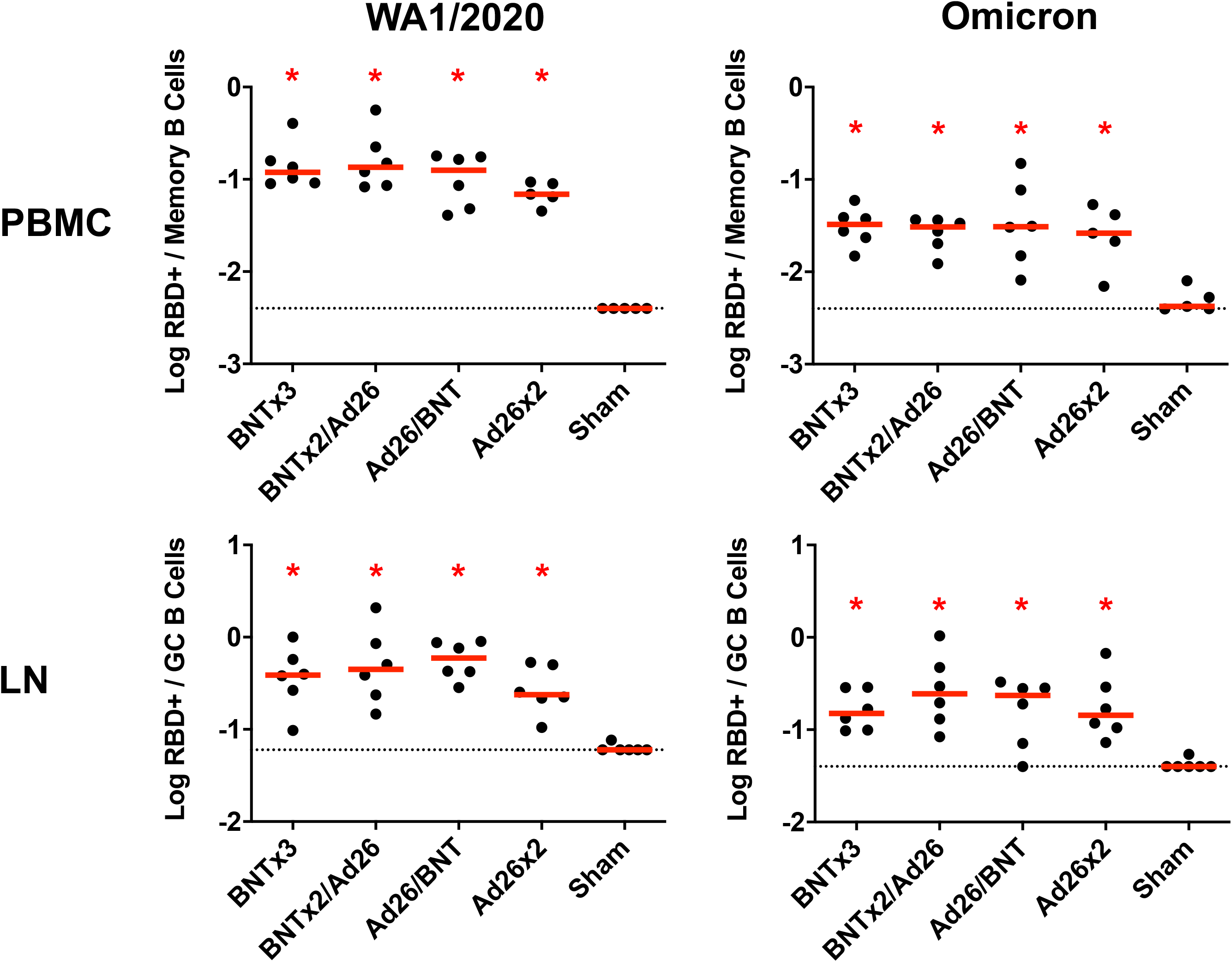
RBD-specific B cell responses following vaccination. WA1/2020 and Omicron RBD-specific memory B cell responses in peripheral blood mononuclear cells (PBMC) and germinal center B cell responses in lymph nodes (LN) are shown at week 16 (post-boost) following vaccination with BNTx3, BNTx2/Ad26, Ad26/BNT, Ad26×2, or sham (N=30; N=6/group). Dotted lines represent limits of quantitation. Medians (red bars) are shown. Vaccinated groups were compared with the sham controls by two-sided Mann-Whitney tests. *, P<0.05.

**Figure S5.**
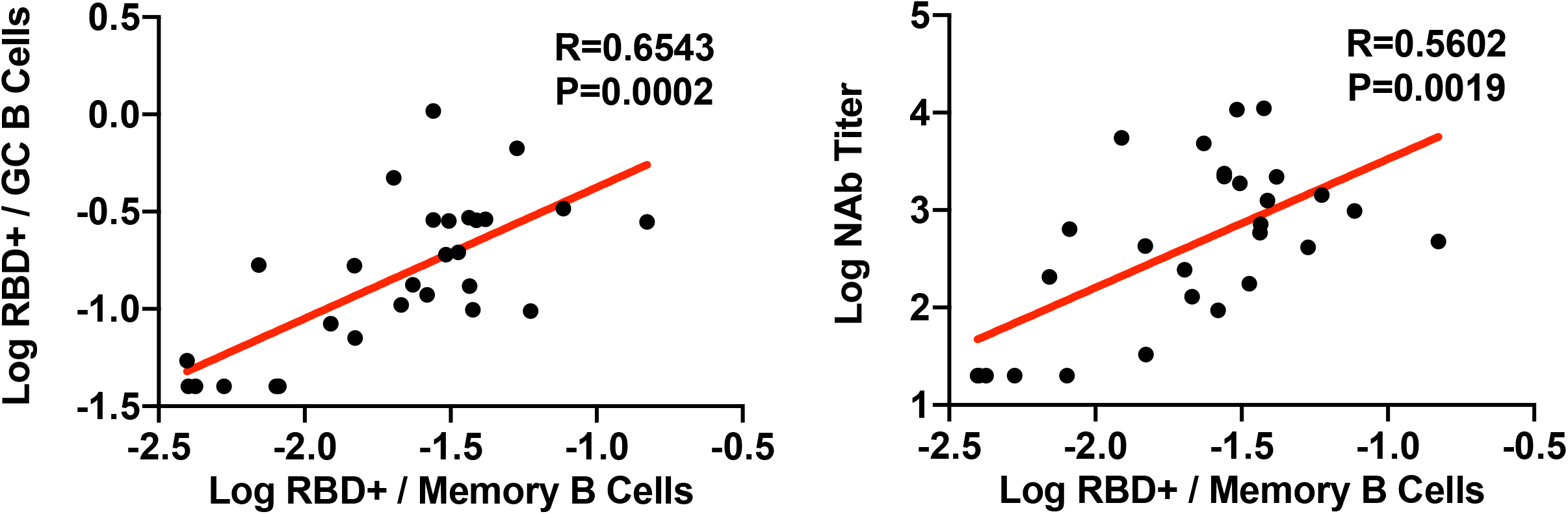
Correlations of RBD-specific B cell responses following vaccination. Correlations of Omicron RBD-specific memory B cell responses in peripheral blood mononuclear cells (PBMC) with Omicron germinal center B cell responses in lymph nodes (LN) (left) and Omicron serum NAb titers are shown at week 16 (post-boost) (right) following vaccination with BNTx3, BNTx2/Ad26, Ad26/BNT, Ad26×2, or sham (N=30; N=6/group). Correlations were assessed by two-sided Spearman rank-correlation tests. R and P values and a regression line of best fit are shown.

**Figure S6.**
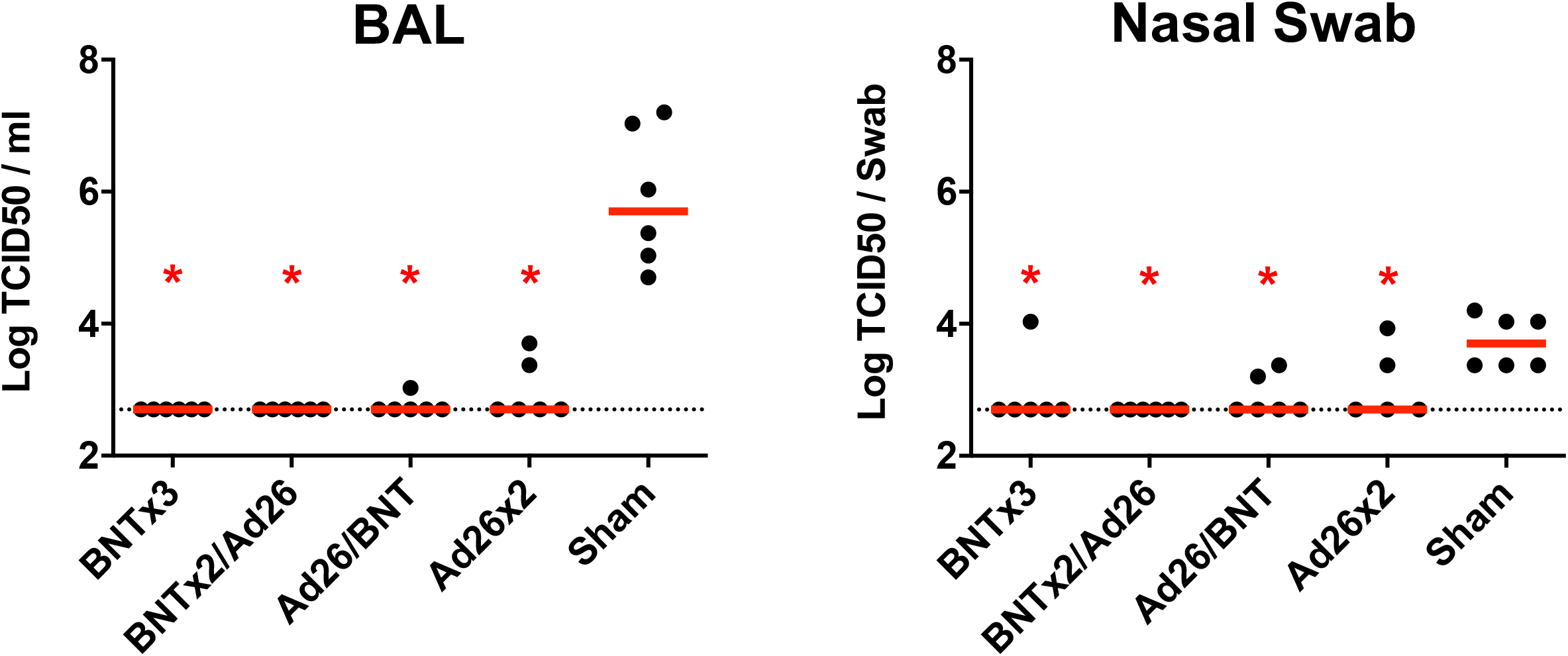
Comparison of day 2 TCID50 titers. Log TCID50/ml in bronchoalveolar lavage (BAL) on day 2 following SARS-CoV-2 Omicron challenge. Dotted lines represent limits of quantitation. Medians (red bars) are shown. Vaccinated groups were compared with the sham controls by two-sided Mann-Whitney tests. *, P<0.05.

**Figure S7.**
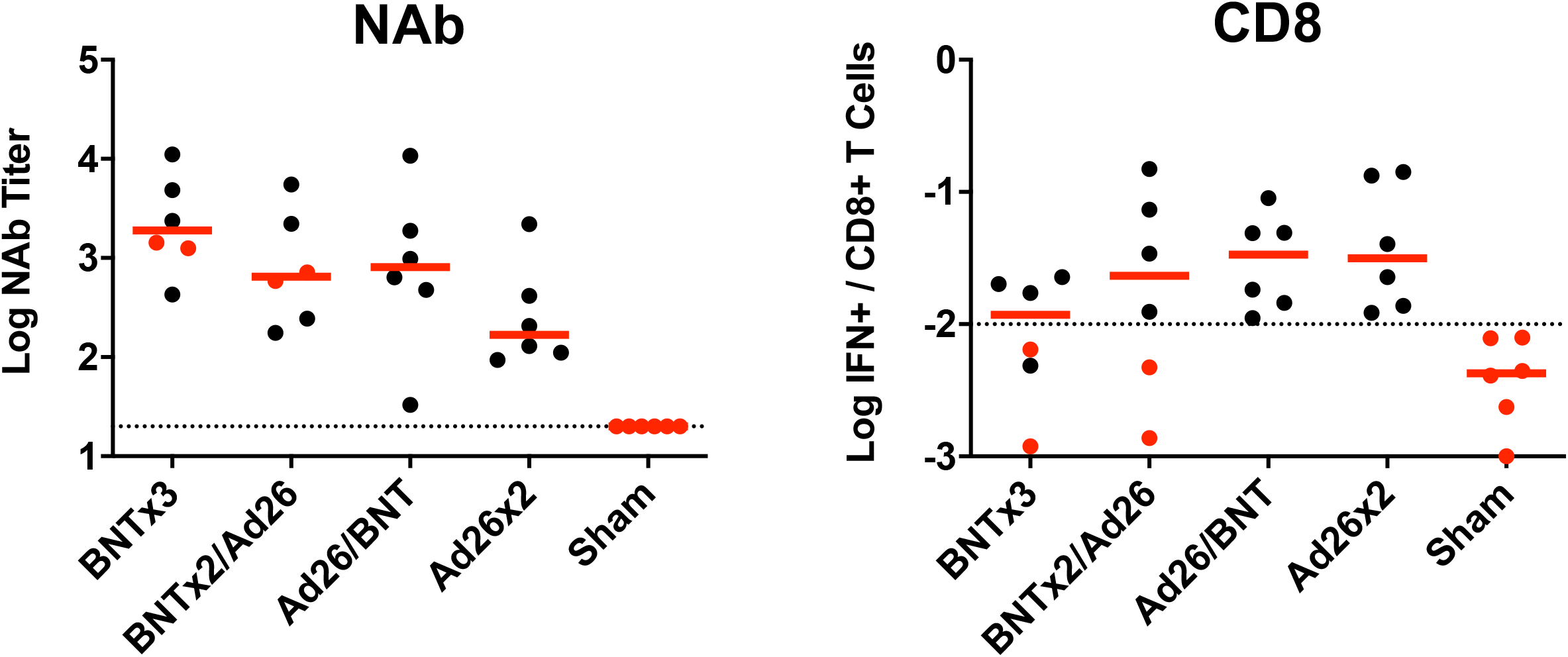
Omicron-specific NAb and CD8+ T cell responses following the boost immunization. Data are extracted from Figures 1, **2**. The 4 animals that failed to show virologic control in NS are highlighted in red (2 in the BNTx3 group, 2 in the BNTx2/Ad26 group).

**Figure S8.**
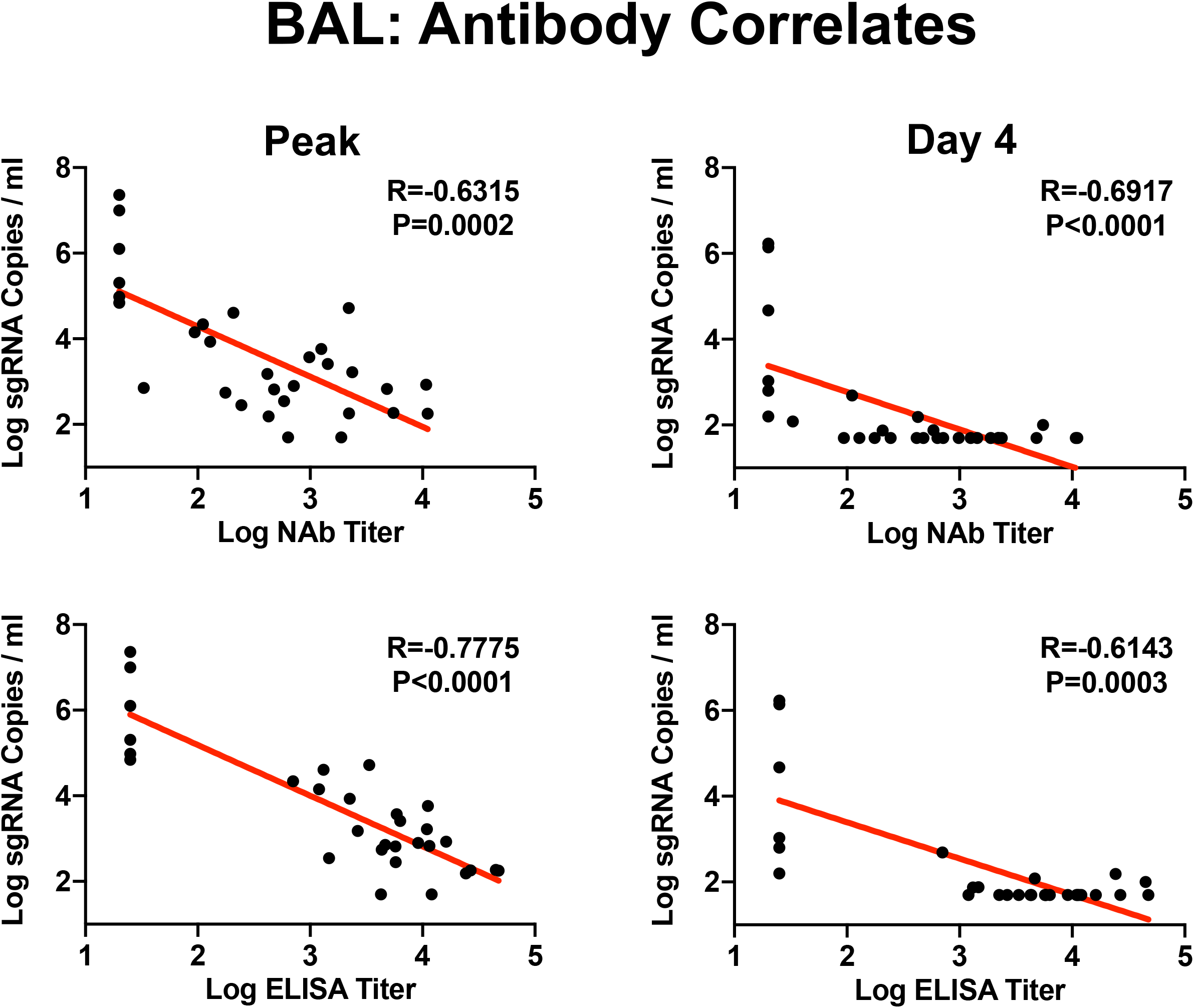
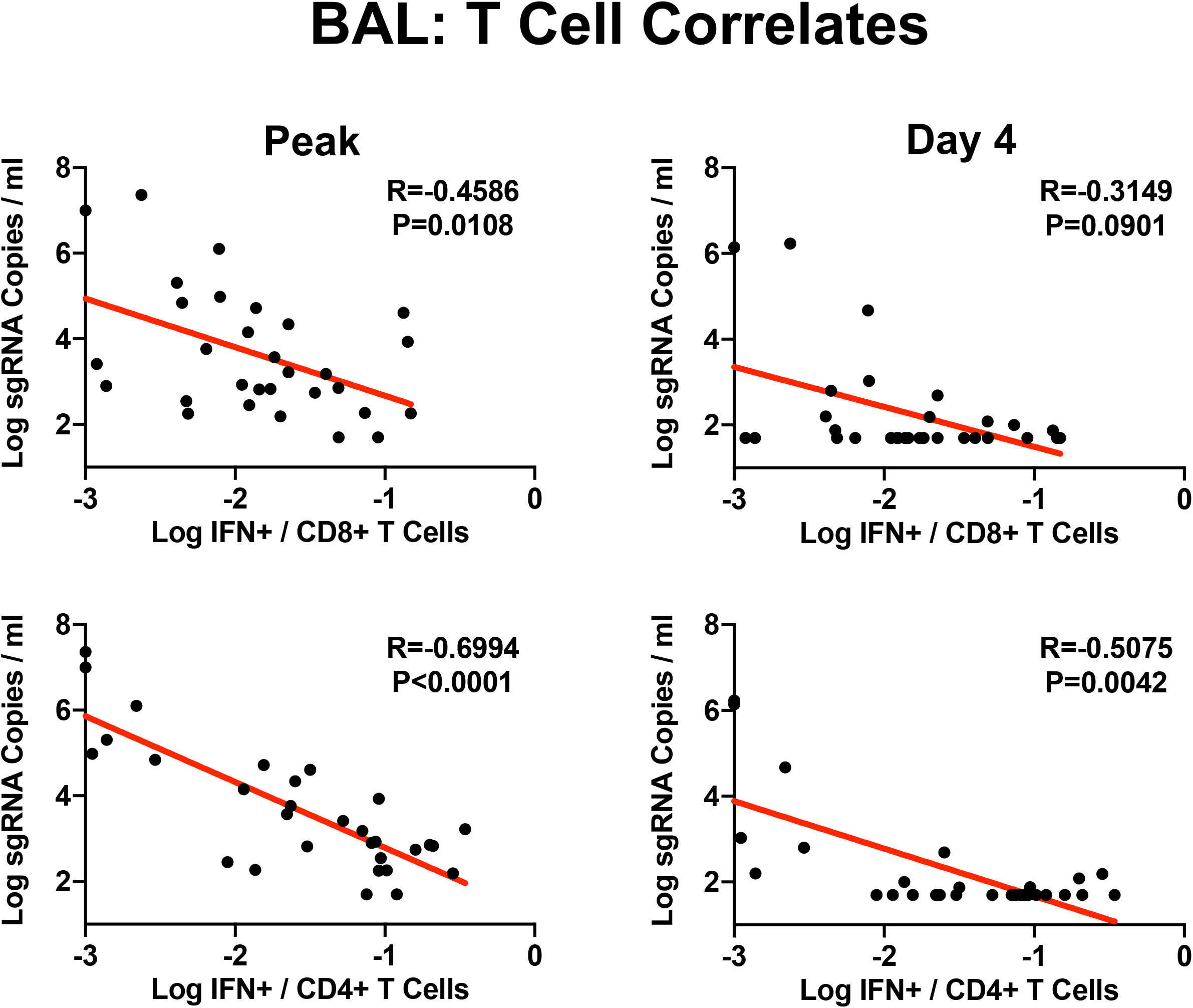
Correlates of protection in BAL. Correlations of week 18 NAb and ELISA titers (**A**) and week 16 CD8+ and CD4+ T cell responses (**B**) with peak and day 4 sgRNA copies/ml in BAL are shown. Correlations were assessed by two-sided Spearman rank-correlation tests. R and P values and a regression line of best fit are shown.

**Figure S9.**
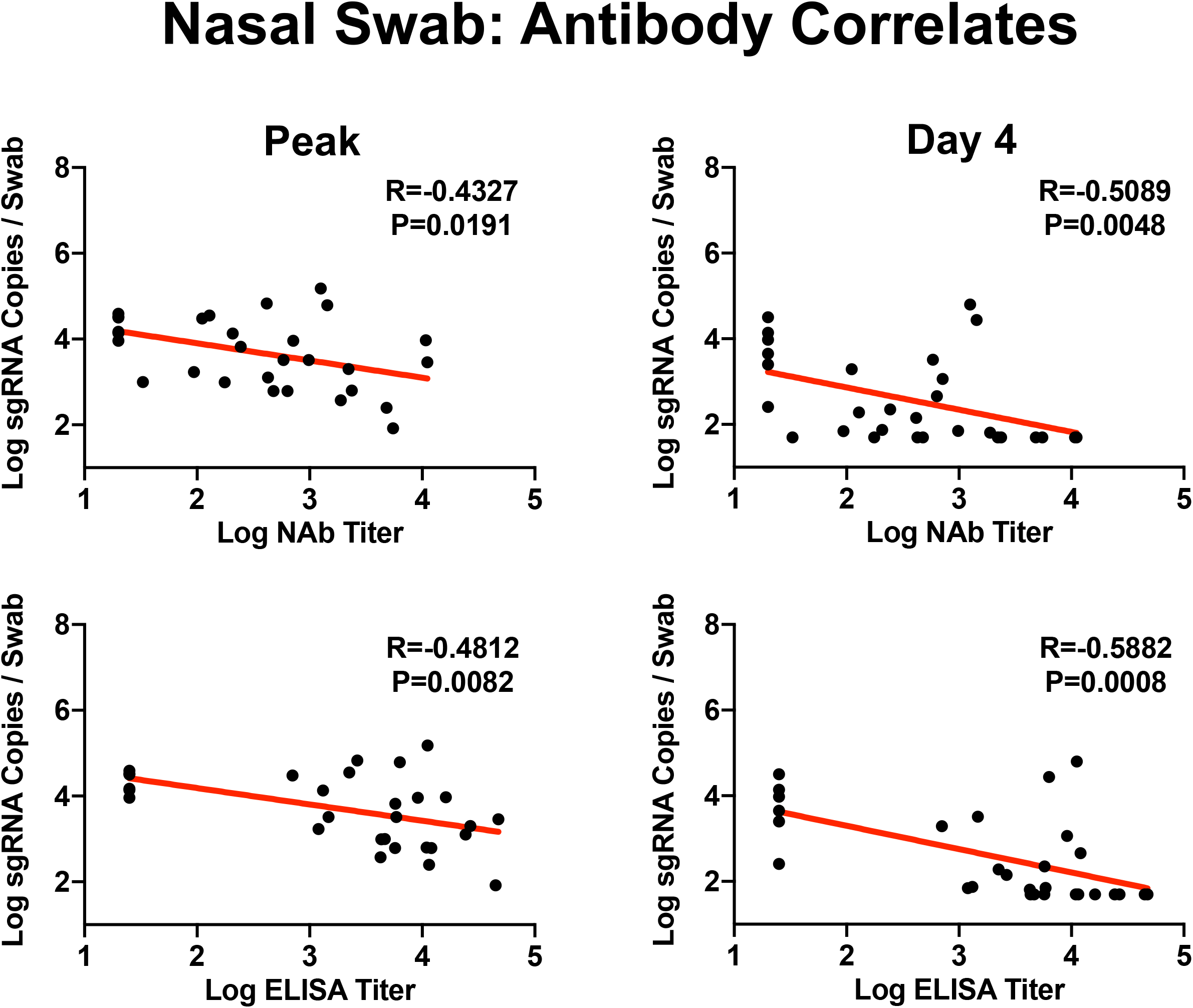
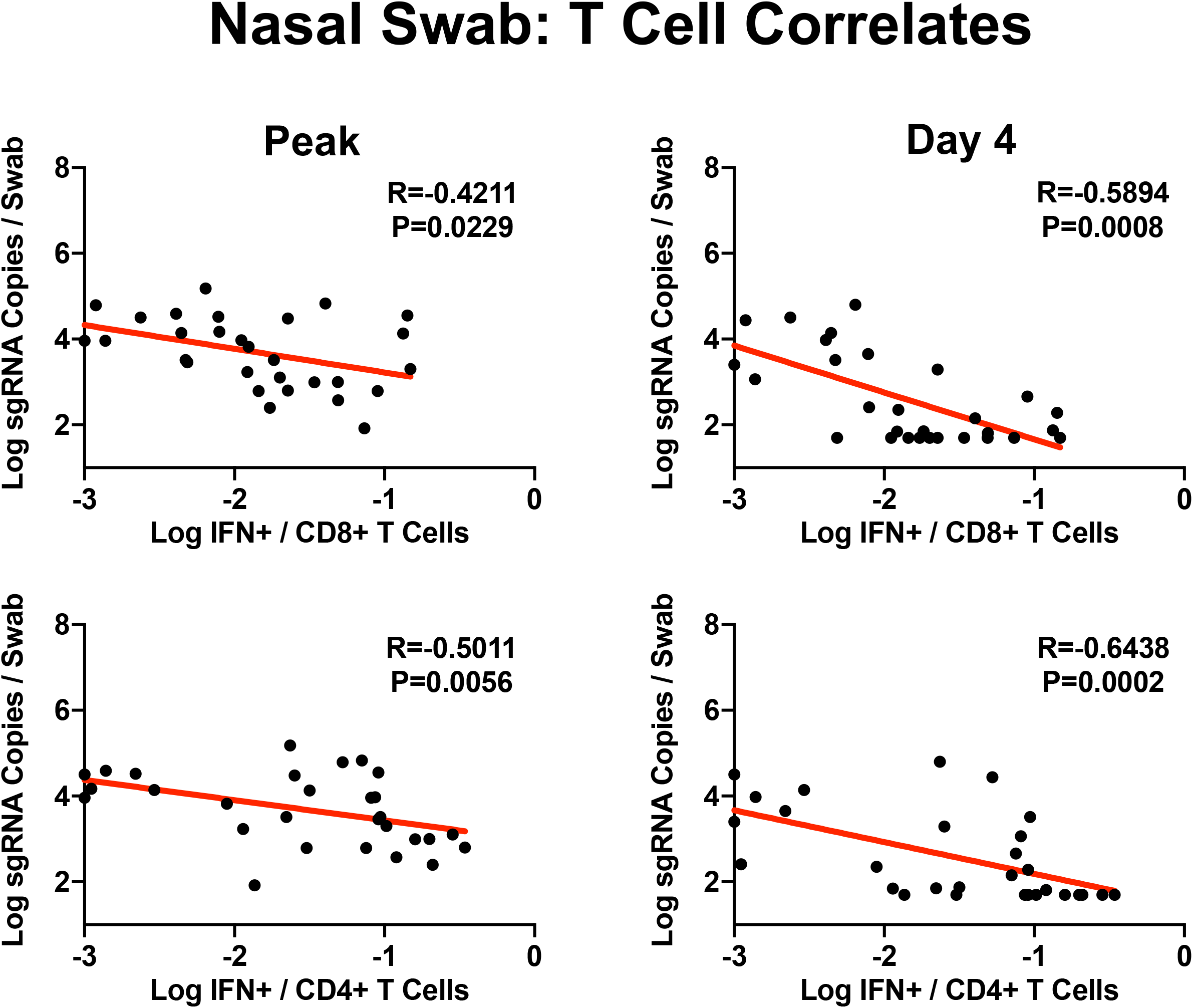
Correlates of protection in NS. Correlations of week 18 NAb and ELISA titers (**A**) and week 16 CD8+ and CD4+ T cell responses (**B**) with peak and day 4 sgRNA copies/swab in NS are shown. Correlations were assessed by two-sided Spearman rank-correlation tests. R and P values and a regression line of best fit are shown.

**Figure S10.**
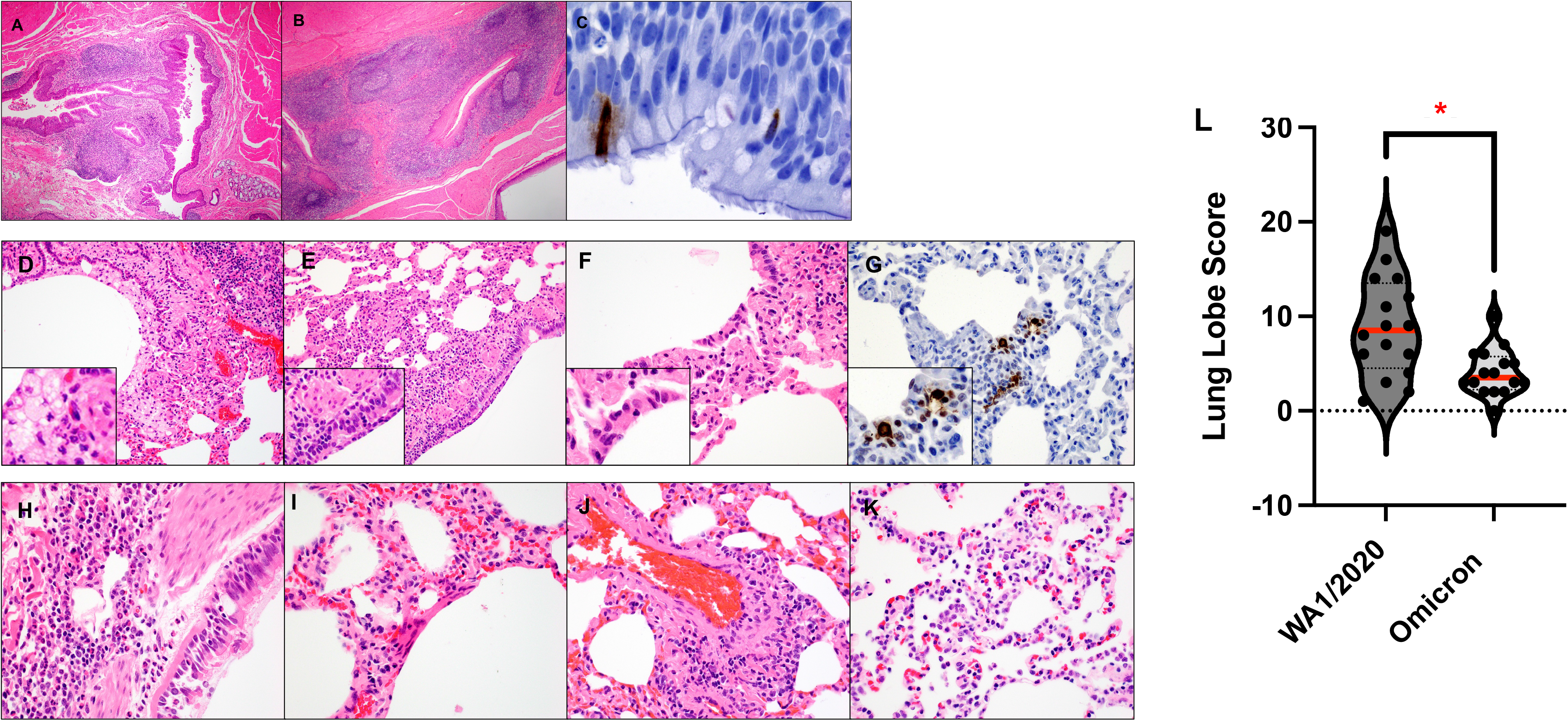
Histopathology and immunohistochemistry of Omicron infection. (**A-C**) Pharynx and (**D-K**) lungs from macaques on day 2 following Omicron infection demonstrated lymphoid hyperplasia of the pharynx (**A**, **B**), SARS-N positive ciliated epithelial cells in the pharynx (**C**), foamy macrophages and degenerating neutrophils in bronchiole lumen (**D**), cellular necrotic debris adhering to bronchiolar ciliated epithelium (**E**), alveolar syncytia (**F**), SARS-N positive ciliated epithelial cells in the pulmonary interstitium (**G**), neutrophilic bronchitis (**H**), hyaline membranes (**I**), endothelialitis (**J**), and type II pneumocyte hyperplasia (**K**). Scoring involved assessment of the following lesions: interstitial inflammation and septal thickening, interstitial infiltrate (eosinophils), interstitial infiltrate (neutrophils), hyaline membranes, interstitial fibrosis, alveolar infiltrate (macrophages), bronchoalveolar infiltrate (neutrophils), epithelial syncytia, type II pneumocyte hyperplasia, bronchi infiltrate (macrophages), bronchi infiltrate (neutrophils), bronchi (hyperplasia of bronchus-associated lymphoid tissue), bronchiolar or peribronchiolar infiltrate (mononuclear cells), perivascular infiltrate (mononuclear cells) and endothelialitis. Each feature assessed was assigned a score of: 0, no substantial findings; 1, minimal; 2, mild; 3, moderate; 4, moderate to severe; 5, marked or severe. Scores were added for all lesions across all lung lobes for each macaque, for a maximum possible score of 600 for each macaque. (**L**) Summary of lung pathology scores from SARS-CoV-2 WA1/2020 and Omicron infected macaques. Medians (red bars) are shown. Dotted line represents no pathology. Vaccinated groups were compared with the sham controls by two-sided Mann-Whitney tests. *, P<0.05.

## REFERENCES

1. Cele, S. et al. Omicron extensively but incompletely escapes Pfizer BNT162b2 neutralization. Nature, doi:10.1038/s41586-021-04387-1 (2021).

2. Liu, L. et al. Striking Antibody Evasion Manifested by the Omicron Variant of SARS-CoV-2. Nature, doi:10.1038/s41586-021-04388-0 (2021).

3. Carreno, J. M. et al. Activity of convalescent and vaccine serum against SARS-CoV-2 Omicron. Nature, doi:10.1038/s41586-022-04399-5 (2021).

4. Schmidt, F. et al. Plasma Neutralization of the SARS-CoV-2 Omicron Variant. N Engl J Med, doi:10.1056/NEJMc2119641 (2021).

5. Nemet, I. et al. Third BNT162b2 Vaccination Neutralization of SARS-CoV-2 Omicron Infection. N Engl J Med, doi:10.1056/NEJMc2119358 (2021).

6. Liu, J. et al. Vaccines Elicit Highly Conserved Cellular Immunity to SARS-CoV-2 Omicron. Nature, doi:10.1038/s41586-022-04465-y (2022).

7. Tarke, A. et al. SARS-CoV-2 vaccination induces immunological T cell memory able to cross-recognize variants from Alpha to Omicron. Cell, doi:https://doi.org/10.1016/j.cell.2022.01.015 (2022).

8. Keeton, R. et al. T cell responses to SARS-CoV-2 spike cross-recognize Omicron. Nature, doi:10.1038/s41586-022-04460-3 (2022).

9. Polack, F. P. et al. Safety and Efficacy of the BNT162b2 mRNA Covid-19 Vaccine. N Engl J Med, doi:10.1056/NEJMoa2034577 (2020).

10. Sadoff, J. et al. Safety and Efficacy of Single-Dose Ad26.COV2.S Vaccine against Covid-19. N Engl J Med 384, 2187–2201, doi:10.1056/NEJMoa2101544 (2021).

11. Collie, S., Champion, J., Moultrie, H., Bekker, L. G. & Gray, G. Effectiveness of BNT162b2 Vaccine against Omicron Variant in South Africa. N Engl J Med, doi:10.1056/NEJMc2119270 (2021).

12. Gray, G. E. et al. Vaccine effectiveness against hospital admission in South African health care workers who received a homologous booster of Ad26.COV2 during an Omicron COVID19 wave: Preliminary Results of the Sisonke 2 Study. medRxiv, 2021.2012.2028.21268436, doi:10.1101/2021.12.28.21268436 (2021).

13. Yu, J. et al. Deletion of the SARS-CoV-2 Spike Cytoplasmic Tail Increases Infectivity in Pseudovirus Neutralization Assays. J Virol, doi:10.1128/JVI.00044-21 (2021).

14. Jacob-Dolan, C. et al. Coronavirus-Specific Antibody Cross Reactivity in Rhesus Macaques Following SARS-CoV-2 Vaccination and Infection. J Virol, doi:10.1128/JVI.00117-21 (2021).

15. He, X. et al. Low-dose Ad26.COV2.S protection against SARS-CoV-2 challenge in rhesus macaques. Cell 184, 3467–3473 e3411, doi:10.1016/j.cell.2021.05.040 (2021).

16. Chandrashekar, A. et al. SARS-CoV-2 infection protects against rechallenge in rhesus macaques. Science 369, 812–817, doi:10.1126/science.abc4776 (2020).

17. Collier, A. Y. et al. Differential Kinetics of Immune Responses Elicited by Covid-19 Vaccines. N Engl J Med 385, 2010–2012, doi:10.1056/NEJMc2115596 (2021).

18. Falsey, A. R. et al. SARS-CoV-2 Neutralization with BNT162b2 Vaccine Dose 3. N Engl J Med, doi:10.1056/NEJMc2113468 (2021).

19. Atmar, R. L. et al. Homologous and Heterologous Covid-19 Booster Vaccinations. N Engl J Med, doi:10.1056/NEJMoa2116414 (2022).

20. Munro, A. P. S. et al. Safety and immunogenicity of seven COVID-19 vaccines as a third dose (booster) following two doses of ChAdOx1 nCov-19 or BNT162b2 in the UK (COV-BOOST): a blinded, multicentre, randomised, controlled, phase 2 trial. Lancet 398, 2258–2276, doi:10.1016/S0140-6736(21)02717-3 (2021).

21. Alter, G. et al. Immunogenicity of Ad26.COV2.S vaccine against SARS-CoV-2 variants in humans. Nature 596, 268–272, doi:10.1038/s41586-021-03681-2 (2021).

22. Dagotto, G. et al. Comparison of Subgenomic and Total RNA in SARS-CoV-2 Challenged Rhesus Macaques. J Virol, doi:10.1128/JVI.02370-20 (2021).

23. Wolfel, R. et al. Virological assessment of hospitalized patients with COVID-2019. Nature, doi:10.1038/s41586-020-2196-x (2020).

24. Gilbert, P. B. et al. Immune correlates analysis of the mRNA-1273 COVID-19 vaccine efficacy clinical trial. Science 375, 43–50, doi:10.1126/science.abm3425 (2022).

25. Feng, S. et al. Correlates of protection against symptomatic and asymptomatic SARS-CoV-2 infection. Nat Med 27, 2032–2040, doi:10.1038/s41591-021-01540-1 (2021).

26. Mercado, N. B. et al. Single-shot Ad26 vaccine protects against SARS-CoV-2 in rhesus macaques. Nature 586, 583–588, doi:10.1038/s41586-020-2607-z (2020).

27. Yu, J. et al. DNA vaccine protection against SARS-CoV-2 in rhesus macaques. Science 369, 806–811, doi:10.1126/science.abc6284 (2020).

28. Yu, J. et al. Protective efficacy of Ad26.COV2.S against SARS-CoV-2 B.1.351 in macaques. Nature 596, 423–427, doi:10.1038/s41586-021-03732-8 (2021).

29. Chandrashekar, A. et al. Prior infection with SARS-CoV-2 WA1/2020 partially protects rhesus macaques against reinfection with B.1.1.7 and B.1.351 variants. Sci Transl Med 13, eabj2641, doi:10.1126/scitranslmed.abj2641 (2021).

30. McMahan, K. et al. Correlates of protection against SARS-CoV-2 in rhesus macaques. Nature 590, 630–634, doi:10.1038/s41586-020-03041-6 (2021).

